# Kinesin-8-specific loop-2 controls the dual activities of the motor domain according to tubulin protofilament shape

**DOI:** 10.1101/2022.02.28.480783

**Authors:** Byron Hunter, Matthieu P.M.H. Benoit, Ana B. Asenjo, Caitlin Doubleday, Daria Trofimova, Hernando Sosa, John S. Allingham

## Abstract

Kinesin-8s are dual-activity motor proteins that can move processively on microtubules and depolymerize microtubule plus-ends, but their mechanism of combining these distinct activities remains unclear. We addressed this by obtaining cryo-EM structures (2.6-3.9 Å) of *Candida albicans* Kip3 in different catalytic states on the microtubule lattice and on a curved microtubule end mimic, as well as a microtubule-unbound *Ca*Kip3-ADP crystal structure (2.0 Å). Together with biochemical analyses of *Ca*Kip3 and kinesin-1 mutants, we define a model that explains the kinesin-8 mechanism. The microtubule depolymerization activity originates in conformational changes of the kinesin-8 motor core that are amplified by its dynamic loop-2. On curved microtubule ends, loop-1 assists depolymerization by inserting into preceding motor domains, forming head-to-tail arrays of kinesin-8s that complement loop-2 contacts with curved tubulin. On straight tubulin protofilaments in the microtubule lattice, extended loop-2-tubulin contacts inhibit conformational changes in the motor core, but in the ADP-Pi state these contacts are relaxed, allowing neck-linker docking for motility. These tubulin shape-induced alternations between pro-microtubule-depolymerization and pro-motility kinesin states, regulated by loop-2, are the key to the dual activity of kinesin-8 motors.

---

Members of the kinesin-8 family are unique, as they often integrate the two enzymatic activities of processive microtubule plus end-directed motility and microtubule depolymerization into a single ~ 40 kDa motor domain^1-4^. In most eukaryotes in which kinesin-8s have been studied, these activities allow them to localize to the ends of cytoplasmic or spindle microtubules, where they regulate mitotic spindle size, spindle position, and chromosomal congression^5-11^. A lack of high-resolution structures capturing a kinesin-8 at each major step of its motile and microtubule depolymerization cycles has limited our understanding of the relationship between these two catalytic cycles and the structural elements of kinesin-8 that control them.

At this time, our knowledge of the catalytic cycles of kinesins is limited to motile kinesins that are incapable of microtubule depolymerization, and to microtubule-depolymerizing kinesins that are non-motile. In motile kinesins, such as kine-sin-1 and kinesin-3, microtubule lattice binding by one of their two motor domains opens the nucleotide-binding pocket, allowing ADP release^12-14^. Subsequent entry of ATP initiates closing of the nucleotide-binding pocket and docking of a short peptide, known as the neck-linker, along the side of the motor domain. Neck-linker docking in the microtubule-bound motor domain propels the second, microtubule-unbound, motor domain towards the microtubule plus end, enabling a single stepping event. In the non-motile microtubule-depolymerizing kinesin-13s, which move on microtubules by diffusion^15^, ATP-binding does not induce closure of the nucleotide-binding pocket on the microtubule lattice^16^. Only when kinesin-13 is bound to curved αβ-tubulin protofilaments found at microtubule ends does the nucleotide-binding pocket close in the presence of ATP^16,17^. In this case, closing the nucleotide-binding pocket allows kinesin-13-specific structural elements to promote further tubulin bending, leading to microtubule depolymerization^18,19^. The motility and depolymerization cycles in purely motile and purely depolymerizing kinesins are thus very different. The features of the kinesin-8 cycle that allow their motors to combine both activities remain unknown.

An important gap in our understanding of the kinesin-8 depolymerization mechanism is the lack of a kinesin-8 structure bound to curved tubulin. Functional studies of *Saccharomyces cerevisiae* kinesin-8 Kip3 (*Sc*Kip3) have proposed that upon reaching the curved conformation of tubulin at the plus end, the motor domain experiences a conformational switch that suppresses ATP hydrolysis^20^. As a result, prolonged, tight interactions between kinesin-8 and the microtubule plus end stabilize protofilament curvature and promote microtubule depolymerization. The suggested mechanism for this tubulin shape-dependent ATPase switch involves formation of an interaction between loop-11 of the motor domain and an aspartate residue on helix-3 of α-tubulin (Asp118 – yeast tubulin, Asp116 – bovine or porcine tubulin). Subsequent molecular dynamics studies identified an arginine within loop-11 (Arg351 – *Sc*Kip3) as a potential candidate residue that interacts with α-tubulin Asp118/116 in the curved conformation^21^. Studies on the human kinesin-8s KIF18A and KIF19A have also implicated separate regions of the microtubule-binding interface, namely loop-2 and loop-8, as elements that are likely involved in stabilizing curved tubulin protofilaments^22-24^. Ultimately, a structure of a kinesin-8 motor bound to curved tubulin is needed to elucidate how discrete parts of the motor domain contribute to kinesin-8-mediated microtubule depolymerization.

Mechanistically, it is well-established that some kinesin-8s operate cooperatively to induce microtubule depolymerization^2,3,20,25^. Both *in vitro* and *in vivo*, a build-up of motors at the plus-end is required for effective disassembly of microtubule polymers. As longer microtubules accumulate more motors than shorter microtubules, longer microtubules are depolymerized faster – an effect termed length-dependent depolymerization. The structural basis underlying this cooperative length-dependent microtubule depolymerization is not understood. It is also not clear if and how kinesin-8s physically interact to collectively produce forces needed to dissociate tubulin dimers.

To understand the mechanistic basis for the dual-functionality of kinesin-8, we obtained seven cryo-EM structures of *Candida albicans* Kip3 bound to either the microtubule lattice or curved microtubule end mimics, and a microtubule-unbound *Ca*Kip3 X-ray structure. We also performed biochemical experiments on *Ca*Kip3 and human kinesin-1 constructs with their loop-2 region swapped or loop-1 region truncated (**Supplementary Fig. 1**). On straight protofilaments within the central microtubule lattice, we observe that loop-2 forms extensive contacts with α-tubulin that temporarily restrict the ATP-bound motor in a non-motile, open conformation. Only in the post-ATP-hydrolysis state of microtubule-bound *Ca*Kip3 are the loop-2-tubulin interactions relaxed enough to allow nucleotide-binding pocket closure and neck-linker docking, producing a pro-motility state. On curved protofilaments, we observe that loop-2 retracts and promotes closing of the nucleotide-binding pocket around ATP, while loop-1 inserts into the preceding motor domain, forming a pro-depolymerization state. When we replaced *Ca*Kip3’s loop-2 with the short loop-2 of kinesin-1, and truncated loop-1, the microtubule depolymerization activity of *Ca*Kip3 decreased substantially. Alternatively, swapping loop-2 from kinesin-1 into *Ca*Kip3 increased the microtubule-stimulated ATPase and microtubule-gliding speed of *Ca*Kip3. These findings show that the kinesin-8-specific extended loop-2 is a central element that coordinates the motility and depolymerase activities of kinesin-8 in accord with the shape of tubulin it binds and that *Ca*Kip3’s microtubule depolymerization activity involves loop-2-amplified conformational changes of its motor domain and inter-motor domain contacts through loop-1. We also show that the unconventional conformations of *Ca*Kip3’s ATPase cycle on microtubules, modulated by loop-2, bear several similarities to kinesin-13s^16^.

## Results

### Structures of CaKip3 reveal a unique ATPase cycle

To understand the molecular mechanisms of kinesin-8 motility and microtubule depolymerization, we solved high-resolution structures of the *Ca*Kip3 motor domain at key intermediates in its motile cycle, as well as its structure on curved αβ-tubulin rings that mimic protofilaments at a depolymerizing microtubule plus end (**Fig. 1**). To visualize the microtubule-unbound ADP state of *Ca*Kip3, we determined a 2.0 Å X-ray crystal structure of a construct containing the motor domain and a slightly truncated neck-linker (*Ca*Kip3-MDN, residues 1-436) (**Fig. 1b, Supplementary Fig. 1b**). Using a longer *Ca*Kip3 construct containing the motor domain, neck-linker, and the first predicted coiled-coil-forming domain (*Ca*Kip3-MDC, residues 1-482) (**Supplementary Fig. 1b**), we obtained 2.6-3.3 Å (**Supplementary Fig. 2**) cryo-electron microscopy (cryo-EM) structures of microtubule-bound *Ca*Kip3 in: (1) a nucleotide-free state (**Fig.1c**), (2) an ATP-like state using the non-hydrolysable ATP analog AMP-PNP (**Fig. 1d**), and (3) an ADP-Pi-like transition state using ADP-AlF_x_ (**Fig. 1e**). To view *Ca*Kip3’s interactions with curved αβ-tubulin protofilaments, we obtained a 3.9 Å cryo-EM structure of *Ca*Kip3-MDC bound to dolastatin-stabilized curved tubulin rings and AMP-PNP (**Fig. 1f, Supplementary Fig. 3**). The data processing and model refinement statistics for the X-ray crystal structure and all cryo-EM structures are reported in **Table 1** and **Table 2**, respectively.

**Fig. 1:**
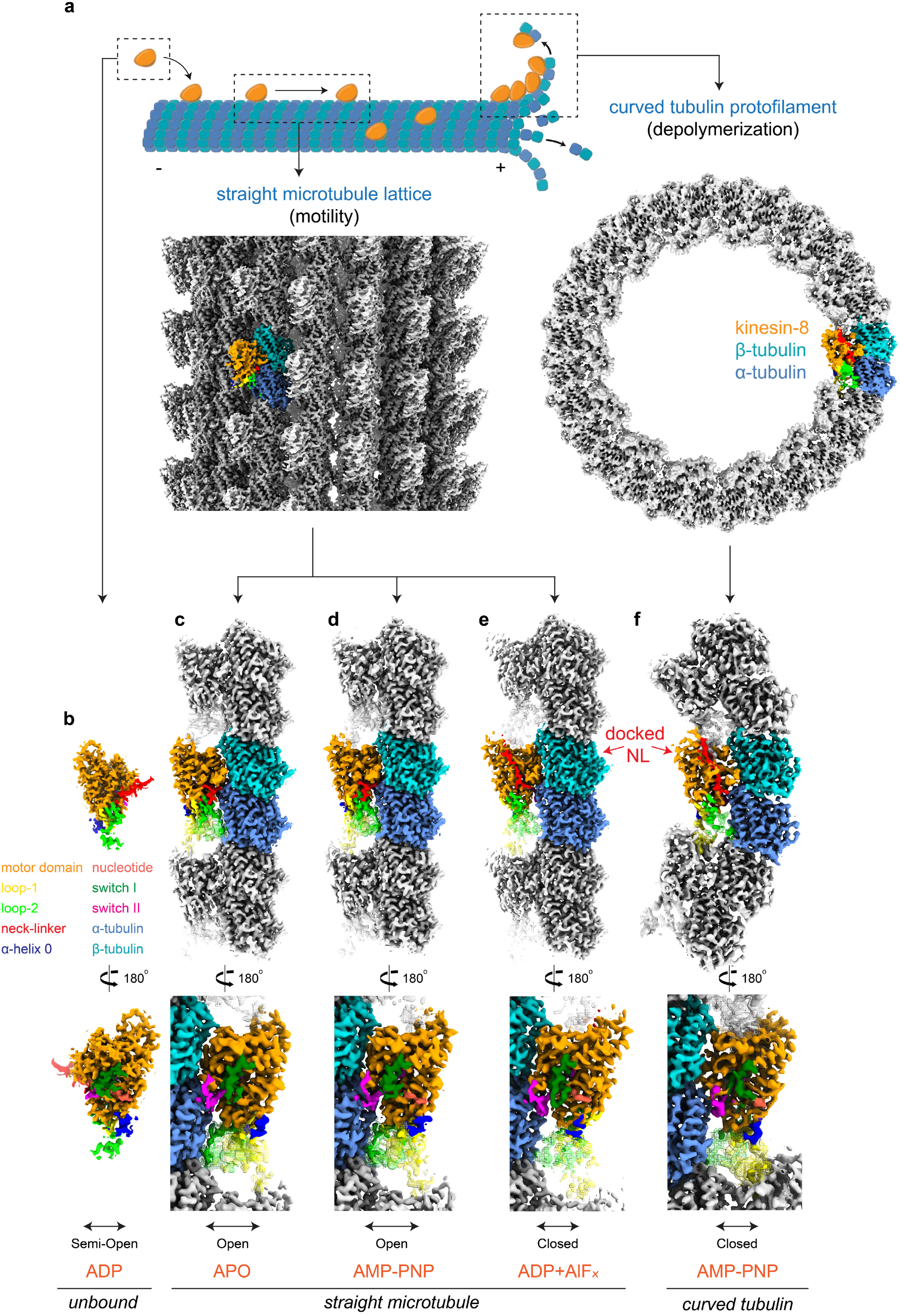
Structures of microtubule-unbound, microtubule-bound, and curved tubulin-bound *Ca*Kip3. **(a)** Top **–** Cartoon representation of catalytic intermediates of *Ca*Kip3’s motility and microtubule depolymerization cycles. Bottom **–** Example of a *Ca*Kip3-decorated microtubule (MT-*Ca*Kip3-MDC-ANP) cryo-EM map and the full *Ca*Kip3-decorated dolastatin-tubulin-ring cryo-EM map. **(b)** X-ray crystallographic density of *Ca*Kip3-MDN in the ADP state. **(c-e)** Cryo-EM maps of microtubule-bound *Ca*Kip3-MDC in the APO, AMP-PNP, and ADP-AlF_x_ nucleotide states. **(f)** Cryo-EM maps of curved tubulin-bound *Ca*Kip3-MDC in the AMP-PNP state. Map surfaces are colored regionally according to the segment of the fitted protein model they enclose: α-tubulin (cornflower blue), β-tubulin (sky blue), kinesin motor core (orange), Switch I loop (forest green), loop-11 of Switch II (magenta), neck-linker (red), nucleotide (tomato), loop-1 (yellow), loop-2 (lime green). The figure was prepared with UCSF ChimeraX^29^.

**Table 1.**
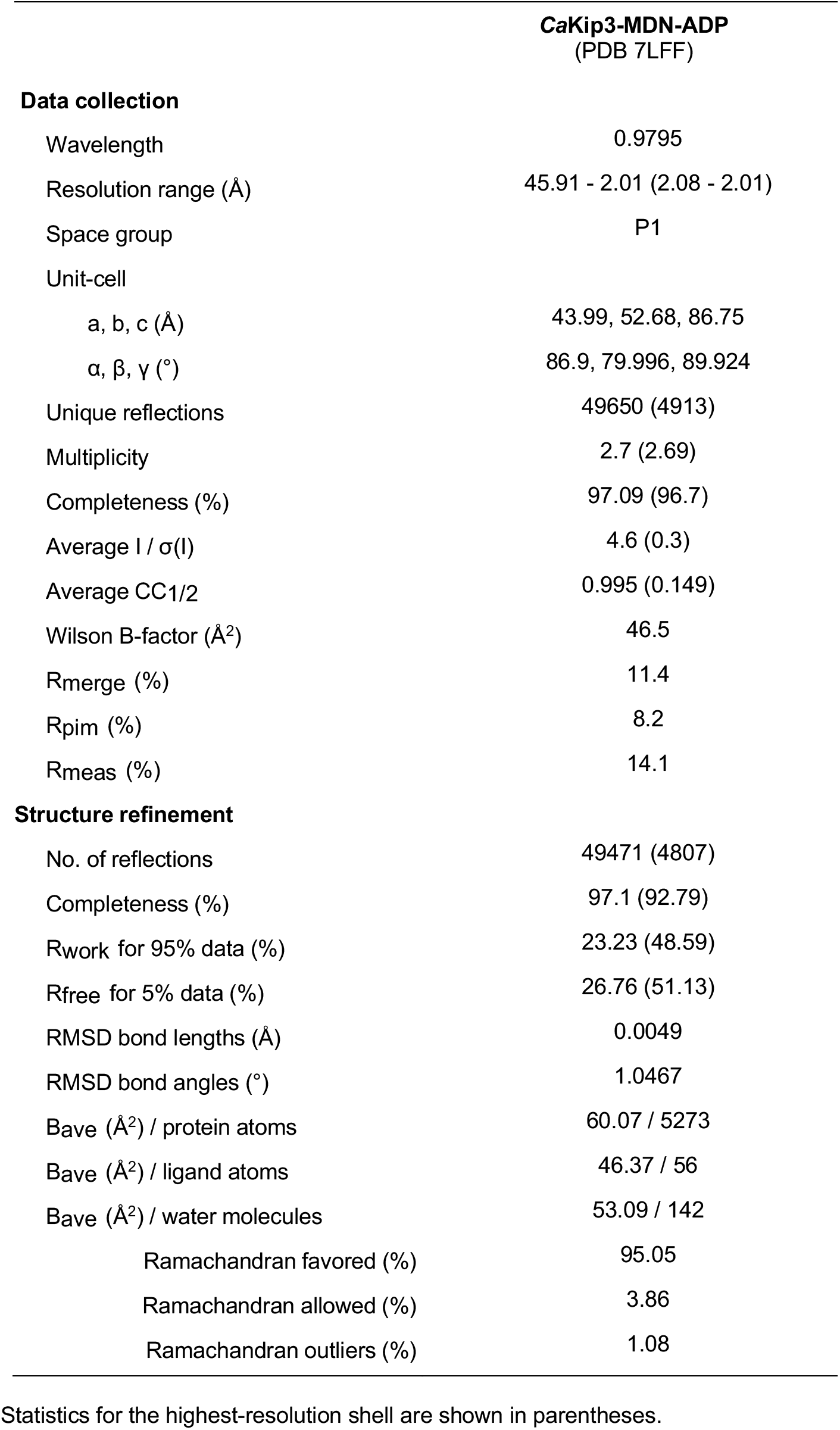
X-ray diffraction data collection, refinement and validation statistics.

**Table 2.** Cryo-EM data collection, refinement and validation statistics (1/2)

|  | **MT-CaKip3-**<br>**MDC**<br>**AMP-PNP**<br>(EMDB-26074)<br>(PDB 7TQX) | **MT-CaKip3-**<br>**MDC**<br>**ADP-AIF<sub>x</sub>**<br>(EMDB-26075)<br>(PDB 7TQY) | **MT-CaKip3-**<br>**MDC**<br>**APO**<br>(EMDB-26076)<br>(PDB 7TQZ) | **MT-CaKip3-**<br>**MDN**<br>**AMP-PNP**<br>(EMDB-26077)<br>(PDB 7TR0) |
| --- | --- | --- | --- | --- |
| **Data collection and processing** |  |  |  |  |
| Magnification (actual) | 60606 | 58893 | 60606 | 60241 |
| Voltage (kV) | 300 | 300 | 300 | 300 |
| Electron exposure (e<sup>-</sup>/Å<sup>2</sup>) | 70.4 | 63.3 | 71.1 | 69.6 |
| Defocus range (μm) <sup>a</sup> | -0.8; -2.1 | -0.8; -1.7 | -0.9; -1.8 | -0.9; -1.7 |
| Pixel size (Å) | 0.825 | 0.849 | 0.825 | 0.830 |
| Symmetry imposed <sup>b</sup> | Helical | Helical | Helical | Helical |
| Rise (Å) | 5.56 | 5.44 | 5.55 | 5.56 |
| Twist (deg) | 168.09 | 168.07 | 168.09 | 168.09 |
| Particle images identified as 15R symmetry (no.) | 12120 | 23783 | 16061 | 77728 |
| Particle images in helical reconstruction (no.) | 12120 | 23783 | 16061 | 77728 |
| Single particles (no.) <sup>c</sup> | 181800 | 356745 | 240915 | 1165920 |
| Single particles used (no.) <sup>d</sup> | 124055 | 255526 | 162955 | 501258 |
| Overall resolution (Å) | 2.8 | 2.6 | 2.7 | 2.7 |
| FSC threshold | 0.143 | 0.143 | 0.143 | 0.143 |
| Kinesin resolution (Å) | 3.1 | 2.9 | 3.0 | 2.9 |
| Tubulin resolution (Å) | 2.8 | 2.6 | 2.7 | 2.7 |
| **Refinement** |  |  |  |  |
| Model composition |  |  |  |  |
| Non-hydrogen atoms | 9824 | 9911 | 9791 | 9836 |
| Protein residues | 1229 | 1241 | 1229 | 1229 |
| Ligands | 6 | 7 | 4 | 5 |
| R.m.s. deviations |  |  |  |  |
| Bond lengths (Å) | 0.0054 | 0.0054 | 0.0052 | 0.0058 |
| Bond angles (°) | 1.14 | 1.17 | 1.12 | 1.17 |
| Validation |  |  |  |  |
| MolProbity score | 1.48 | 1.36 | 1.41 | 1.36 |
| Clashscore | 4.28 | 3.23 | 4.08 | 4.17 |
| Poor rotamers (%) | 0.47 | 1.12 | 0.94 | 0.56 |
| Ramachandran plot |  |  |  |  |
| Favored (%) | 96.07 | 96.66 | 96.64 | 97.13 |
| Allowed (%) | 3.93 | 3.34 | 3.36 | 2.87 |
| Disallowed (%) | 0.00 | 0.00 | 0.00 | 0.00 |

<sup>a</sup> Range of the per-particle defocus estimates. The range corresponds to 90% of the particles used, 5% of the particle defocusses are below and 5% are above this range.
<sup>b</sup> Symmetry imposed during the symmetrical refinement done before the local refinements (rise and twists provided for helical symmetry only).
<sup>c</sup> Total number of particles after symmetry expansion.
<sup>d</sup> Number of asymmetric units, corresponding to a kinesin motor bound to a tubulin dimer, used after 3D classification.

Table 2. continued (2/2)
|  | MT-CaKip3-MDN<sub>L2-KHC</sub><br>AMP-PNP<br>(EMDB-26078)<br>(PDB 7TR1) | MT-CaKip3-MDN<sub>L2-KHC</sub><br>APO<br>(EMDB-26079)<br>(PDB 7TR2) | CT-CaKip3-MDC<br>AMP-PNP<br>(EMDB-26080)<br>(PDB 7TR3) |
| --- | --- | --- | --- |
| **Data collection and processing** |  |  |  |
| Magnification (actual) | 45872 | 58893 | 46168 |
| Voltage (kV) | 300 | 300 | 300 |
| Electron exposure (e<sup>-</sup>/Å<sup>2</sup>) | 64.0 | 64.6 | 64.0 |
| Defocus range (μm) <sup>a</sup> | -0.8; -1.9 | -0.8; -1.7 | -0.9; -3.2 |
| Pixel size (Å) | 1.090 | 0.849 | 1.083 |
| Symmetry imposed <sup>b</sup> | Helical | Helical | C14 |
| Rise (Å) | 5.54 | 5.55 |  |
| Twist (deg) | 168.08 | 168.09 |  |
| Particle images identified as 15R/C14 symmetry (no.) | 37146 | 12219 | 37920 |
| Particle images in helical/C14 reconstruction (no.) | 37146 | 12219 | 37920 |
| Single particles (no.) <sup>c</sup> | 557190 | 183285 | 530936 |
| Single particles used (no.) <sup>d</sup> | 67238 | 98446 | 530936 |
| Overall resolution (Å) | 3.1 | 3.0 | 3.9 |
| FSC threshold | 0.143 | 0.143 | 0.143 |
| Kinesin resolution (Å) | 3.4 | 3.4 | 3.9 |
| Tubulin resolution (Å) | 3.0 | 3.0 | 3.9 |
| **Refinement** |  |  |  |
| Model composition |  |  |  |
| Non-hydrogen atoms | 9708 | 9588 | 10223 |
| Protein residues | 1216 | 1207 | 1288 |
| Ligands | 6 | 4 | 5 |
| R.m.s. deviations |  |  |  |
| Bond lengths (Å) | 0.0055 | 0.0054 | 0.0051 |
| Bond angles (°) | 1.14 | 1.15 | 1.12 |
| Validation |  |  |  |
| MolProbity score | 1.44 | 1.35 | 1.79 |
| Clashscore | 3.39 | 2.96 | 5.36 |
| Poor rotamers (%) | 0.19 | 1.05 | 0.27 |
| Ramachandran plot |  |  |  |
| Favored (%) | 95.61 | 96.33 | 91.58 |
| Allowed (%) | 4.39 | 3.67 | 8.42 |
| Disallowed (%) | 0.00 | 0.00 | 0.00 |

<sup>a</sup> Range of the per-particle defocus estimates. The range corresponds to 90% of the particles used, 5% of the particle defocuses are below and 5% are above this range.
<sup>b</sup> Symmetry imposed during the symmetrical refinement done before the local refinements (rise and twists provided for helical symmetry only).
<sup>c</sup> Total number of particles after symmetry expansion.
<sup>d</sup> Number of asymmetric units, corresponding to a kinesin motor bound to a tubulin dimer, used after 3D classification.

Comparing the different *Ca*Kip3 structures reveals that the conformational changes experienced by the kinesin-8 motor domain core during microtubule binding in relation to its ATP hydrolysis cycle are different from purely motile kinesins. The *Ca*Kip3-MDN-ADP crystal structure has a partially open nucleotide pocket and an undocked neck-linker (**Fig. 1b, Supplementary Fig. 4**). The Switch I region (loop-9) is well defined but angled away from ADP, while the Switch II region (loop-11) that coordinates microtubule sensing and nucleotide binding is mostly disordered. When *Ca*Kip3-MDC binds the microtubule (MT-*Ca*Kip3-MDC-APO), the minus subdomain rotates relative to the plus subdomain, opening the nucleotide-binding pocket further (**Fig. 1c, Fig. 2a**). Here, the full length of loop-11 is visible and interacts with helices 3’, 3, 11’, and 12 of α-tubulin. Unexpectedly, binding of ATP to the MT-*Ca*Kip3-MDC complex, as mimicked by AMP-PNP, fails to induce a ‘pro-motility’ conformational change (i.e., nucleotide-binding pocket closure and neck-linker docking on the microtubule lattice) even though the *Ca*Kip3-MDC construct includes the full neck-linker (**Fig. 1d, Fig. 2b, Supplementary Fig. 6**). The cryo-EM map of the MT-*Ca*Kip3-MDC-ANP complex clearly shows that the nucleotide-binding pocket is open, and the neck-linker is undocked, similar to MT-*Ca*Kip3-MDC-APO (compare **Fig. 1c** and **1d**). This is striking because all microtubule-bound structures of motile kinesins with a full neck-linker that is not pulled backward by a trailing head exhibit a closed nucleotide-binding pocket in the ATP-bound state (as mimicked by AMP-PNP) (**compare Supplementary Fig. 6b, 6c and 6i**)^13,26-28^. In this regard, *Ca*Kip3 is more akin to depolymerizing kinesins than to motile kinesins^16^. Only in the post-ATP-hydrolysis state (MT-*Ca*Kip3-MDC-AAF) does the microtubule-bound *Ca*Kip3 motor domain adopt the pro-motility conformation with a closed nucleotide-binding pocket and docked neck-linker, allowing *Ca*Kip3 to complete its motile cycle (**Fig. 1e, Fig. 2c, Supplementary Fig. 2b**).

**Fig. 2:**
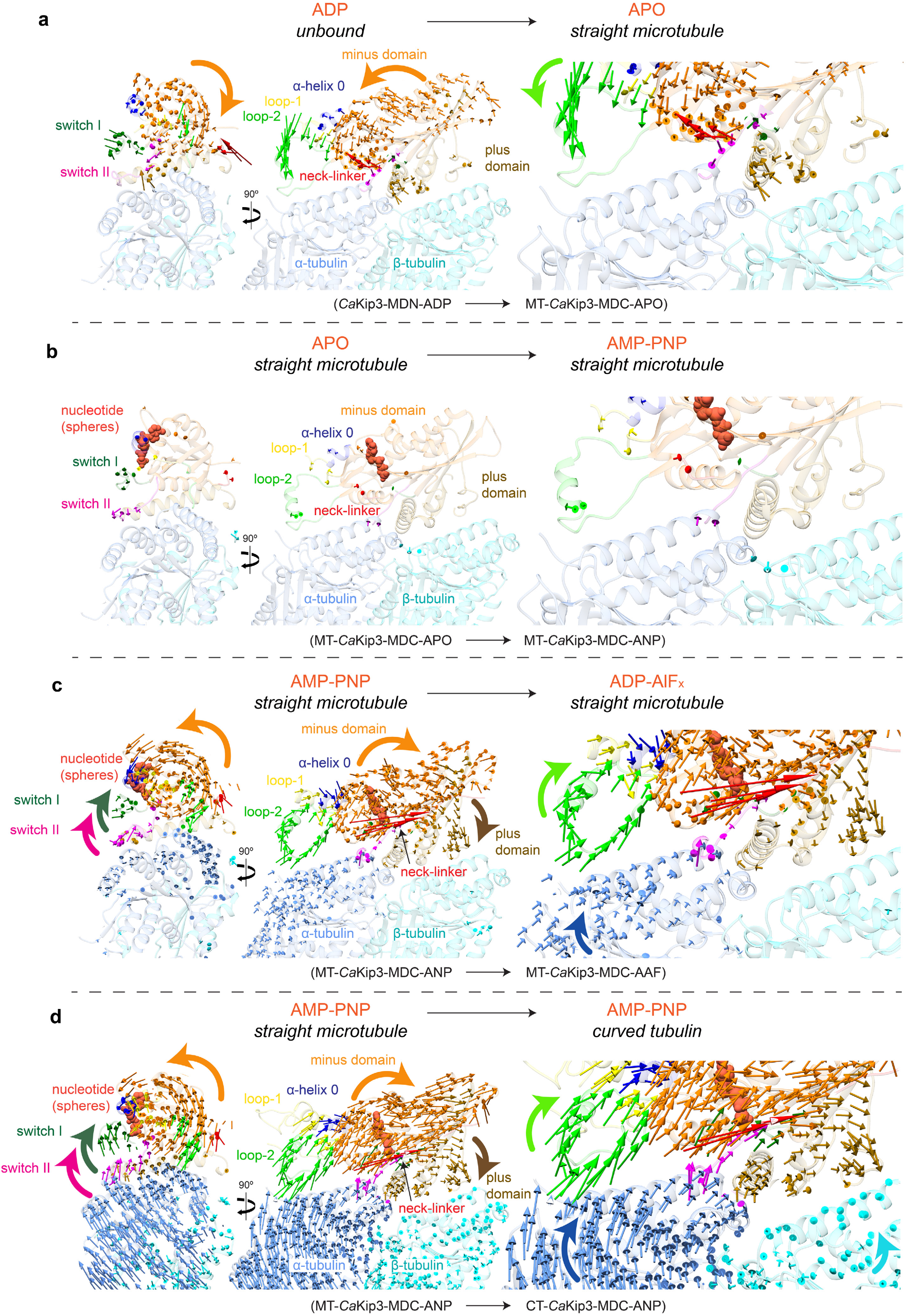
Conformational changes in *Ca*Kip3 and tubulin. **(a)** Displacement vectors for Cα atoms in *Ca*Kip3 when comparing the microtubule-unbound *Ca*Kip3-MDN-ADP structure and the MT-*Ca*Kip3-MDC-APO complex. The models were aligned to microtubule interacting regions H4, loop-11 and loop-8 of the MT-*Ca*Kip3-MDC-APO structure, specifically, residues 249-257 and 350-379. Displacement vectors for Cα atoms in *Ca*Kip3 and tubulin when comparing **(b)** MT-*Ca*Kip3-MDC-APO and MT-*Ca*Kip3-MDC-ANP, **(c)** MT-*Ca*Kip3-MDC-ANP and MT-*Ca*Kip3-MDC-AAF, and **(d)** MT-*Ca*Kip3-MDC-ANP and CT-*Ca*Kip3-MDC-ANP. All structures in **(b-d)** were aligned to the β-tubulin chain of the MT-*Ca*Kip3-MDC-APO complex. Displacement vectors for Cα atoms are colored regionally to match the segment of the protein model that is compared, using the color scheme in **Fig. 1**. Views are from the minus end, down the long axis of the protofilament (left), side view (middle) and close-up of the tubulin interface (right).

Our library of structures defines the subdomain motions associated with nucleotide-binding pocket closure in *Ca*Kip3. When the nucleotide-binding pocket closes in the MT-*Ca*Kip3-MDC-AAF complex, the minus subdomain has a strong rotational component in the plane perpendicular to the microtubule axis (**Fig. 2c**). We observe these same conformational changes in *Ca*Kip3-MDC when it is bound to curved tubulin rings and AMP-PNP (CT-*Ca*Kip3-MDC-ANP) (**Fig. 2d**). When viewed from the microtubule minus end, it is evident that closure of the nucleotide-binding pocket is associated with a counter-clockwise rotation of the minus sub-domain and a clockwise rotation of the Switch I and II regions (**Fig. 2c, d; left panels**). Movement of these subdomains in the plane parallel to the microtubule axis results in a relative displacement of the tubulin-interacting regions to better fit curved tubulin, which is similar overall to what was reported in kinesin-13 (**Supplementary Fig. 6h**)^16^. In the MT-*Ca*Kip3-MDC-AAF and CT-*Ca*Kip3-MDC-ANP complexes, the displacement of tubulin-interacting regions relative to β-tubulin is greatest at the kinesin-8-specific elongated loop-2 that extends the kinesin-tubulin interface (**Fig. 2c, d; right panels**). Prominent as well is movement of the loop-8 lobe toward helix-12 of β-tubulin. Unexpectedly, we also observe a small rotation of α-tubulin relative to β-tubulin in the MT-*Ca*Kip3-MDC-AAF complex, giving a slight outward curvature to each *Ca*Kip3-bound αβ-tubulin subunit in the microtubule lattice when the nucleotide-binding pocket of *Ca*Kip3 is closed (**Fig. 2c; right panels – blue arrows on α-tubulin**).

### Extensive and variable loop-2 contacts with α-tubulin

Collectively, our *Ca*Kip3 structural data shows well-defined map density for the lengthy loop-2 region that extends out of the β1b/β1c strands, toward the minus end of the microtubule. The electron density from the *Ca*Kip3-MDN-ADP crystal allowed us to model residues Phe116 to Ala131 of loop-2, which includes a proline-rich region and a ~10-residue helical sequence that is shared by several kinesin-8s (**Fig. 3a, Supplementary Fig. 1a**). The cryo-EM maps of MT-*Ca*Kip3-MDC-APO and MT-*Ca*Kip3-MDC-ANP show that the section of loop-2 that precedes the β1c strand, which is not resolved in the X-ray structure, becomes ordered on the microtubule while *Ca*Kip3 is in the open conformation (**Fig. 3c, d; left panels**). This section interacts with helix-12 and a loop between helix-7 and the β7 strand of α-tubulin via residues Phe136, Ser139, Arg140, and one of the two visible conformations of the side chain of His144 (**Fig. 3c, d; right panels**). These loop-2-tubulin interactions are more numerous than the loop-2 contacts made by kinesin-13s^16^, which are essential for kinesin-13’s microtubule depolymerization activity^16,18,19,30-32^. Several of the *Ca*Kip3 loop-2-tubulin helix-12 interactions are electrostatic. For example, Arg140 is close enough to α-tubulin to hydrogen bond with Glu420 and Asp424 in helix-12. These contacts support earlier assumptions that loop-2 contributes positively charged residues that associate with the negatively charged surface of α-tubulin^23^. We observe similar loop-2-tubulin contacts, and overall motor domain conformation in the two microtubule-bound AMP-PNP structures, MT-*Ca*Kip3-MDC-ANP and MT-*Ca*Kip3-MDN-ANP (**Fig. 4c**).

**Fig. 3:**
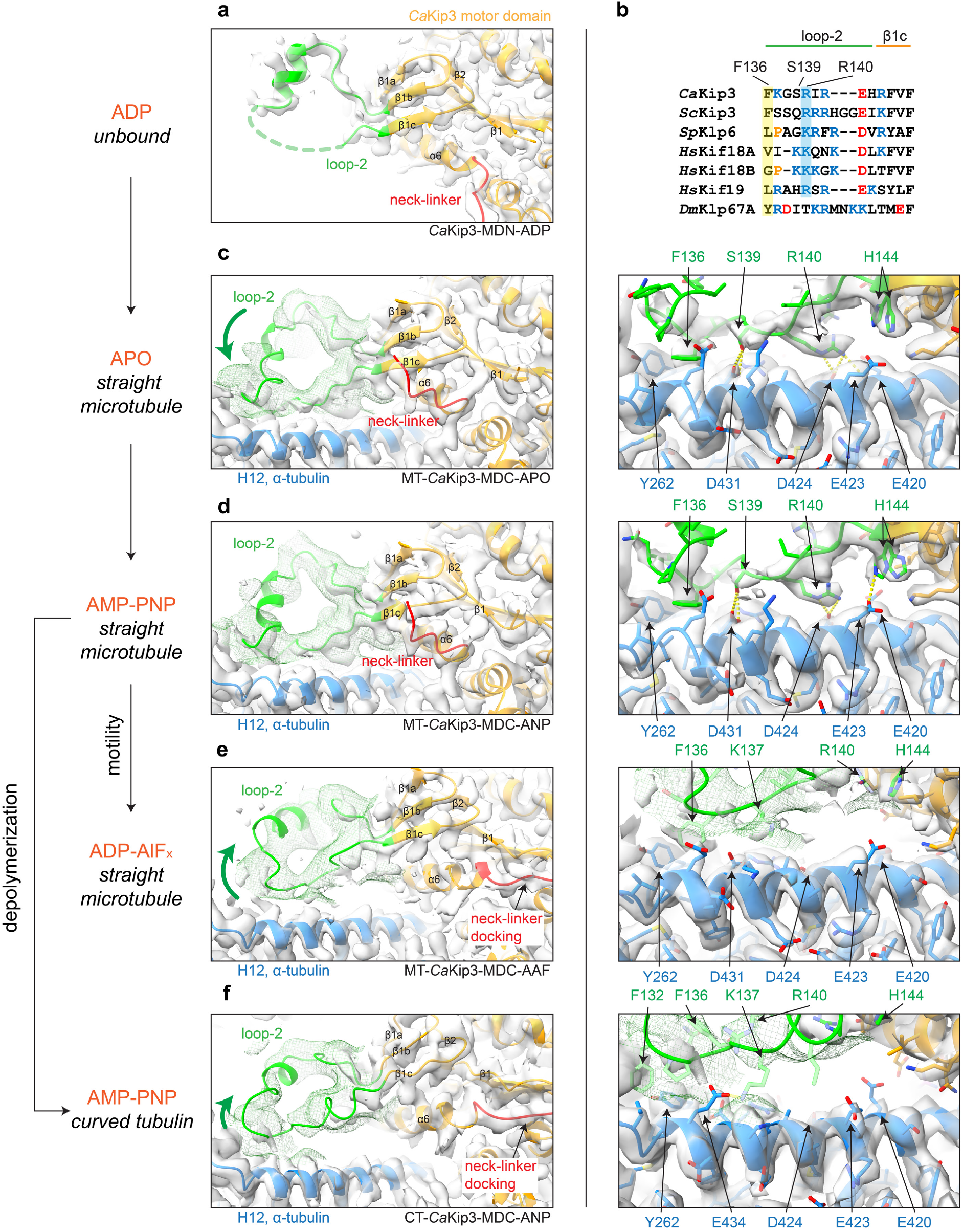
Loop-2 electron and cryo-EM densities and tubulin contacts. **(a)** 2*mF*_obs_ – *dF*_calc_ electron density map (contoured at 1.0*σ*) and cartoon model of loop-2 and the motor domain of the *Ca*Kip3-MDN crystal structure. **(b)** Sequence alignment of the tubulin contact region of *Ca*Kip3’s loop-2 with other kinesin-8s. Conserved tubulin-bonding residues are highlighted. Cryo-EM map and cartoon model of loop-2 of the **(c)** MT-*Ca*Kip3-MDC-APO complex, **(d)** MT-*Ca*Kip3-MDC-ANP complex, **(e)** MT-*Ca*Kip3-MDC-AAF complex, and **(f)** CT-*Ca*Kip3-MDC-ANP complex. Right panels show close-up view of loop-2-tubulin interactions for each complex. Green mesh maps were low-pass filtered at 7 Å (MT-*Ca*Kip3-MDC-AAF at 9 Å) to better display the noisier density of loop-2. Interacting residues are shown in stick representation. Pseudo-bonds between interacting atoms were determined and displayed using the “find clashes and contacts” routine in UCSF ChimeraX and are represented as yellow dashed lines.

**Fig. 4:**
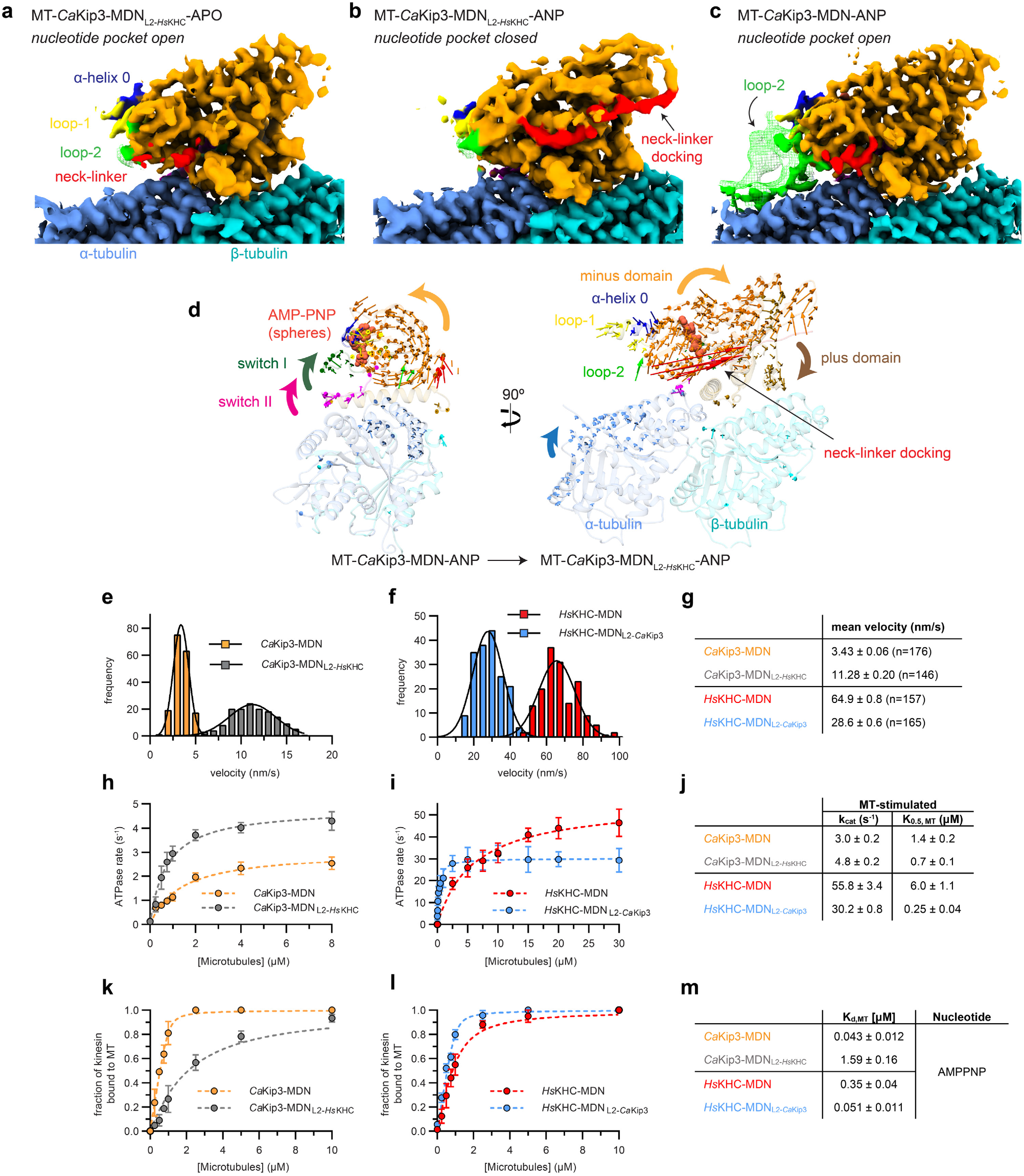
Structures and activity of loop-2 swap mutants on microtubules. Cryo-EM map of **(a)** MT-*Ca*Kip3-MDN_L2-*Hs*KHC_ in the APO state, **(b)** MT-*Ca*Kip3-MDN_L2-*Hs*KHC_ in the AMP-PNP state, and **(c)** MT-*Ca*Kip3-MDN in the AMP-PNP state. **(d)** Displacement vectors for Cα atoms in *Ca*Kip3 and tubulin when comparing the MT-*Ca*Kip3-MDC-ANP and MT-*Ca*Kip3-MDN_L2-*Hs*KHC_-ANP complexes. Structures were aligned to the β-tubulin chain of MT-*Ca*Kip3-MDC-ANP. **(e,f)** Microtubule gliding assay velocity distributions for *Ca*Kip3-MDN, *Ca*Kip3-MDN_L2-*Hs*KHC_, *Hs*KHC-MDN, and *Hs*KHC-MDN_L2-*Ca*Kip3_ using taxol-stabilized microtubules. Microtubules were tracked from 2-3 independent experiments for each protein. **(g)** Summary of motor velocities. Values are presented as mean ± SEM. **(h)** Microtubule-stimulated ATPase kinetics of *Ca*Kip3-MDN and *Ca*Kip3-MDN_L2-*Hs*KHC_ at steady state (n=3). The basal rate was 0.13 ± 0.01 for both motors. **(i)** Microtubule-stimulated ATPase kinetics of *Hs*KHC-MDN and *Hs*KHC-MDN_L2-*Ca*Kip3_ at steady state (n=3). Basal rates were 0.012 ± 0.001 s^-1^ and 0.010 ± 0.001 s^-1^, respectively. **(j)** Summary of the microtubule-stimulated ATPase activities of the motors. Values are presented as mean ± SEM. Data was fit to the Michaelis-Menten equation using GraphPad Prism to obtain *K*_0.5, MT_ and *k*_cat_ values. **(k)** Microtubule-co-sedimentation data for *Ca*Kip3-MDN and *Ca*Kip3-MDN_L2-*Hs*KHC_ (1 μM each) in the presence of 2 mM MgAMP-PNP. **(l)** Microtubule-co-sedimentation data for *Hs*KHC-MDN and *Hs*KHC-MDN_L2-*Ca*Kip3_ (1 μM each) in the presence of 2 mM MgAMP-PNP. Taxol-stabilized microtubules were pelleted by centrifugation to separate the free kinesin and microtubule-bound kinesin. SDS-PAGE and Coomassie brilliant blue staining were used to determine the fraction of microtubule-bound kinesin. **(m)** Microtubule-binding affinities (*K*_d_ values) of the motors were calculated using the quadratic equation given in Materials and Methods. Values are presented as mean ± SEM for three independent experiments.

In the MT-*Ca*Kip3-MDC-AAF and CT-*Ca*Kip3-MDC-ANP complexes, the loop-2 density is much weaker, indicating a more mobile loop structure. We could trace the average path of loop-2 in these complexes by applying a low-pass filter of 9 Å and 7 Å, respectively (**Fig. 3e, f; left panels**). In CT-*Ca*Kip3-MDC-ANP, stronger density is visible along α-tubulin, especially near the loop-2 tip around α-tubulin residue Y262, compared to MT-*Ca*Kip3-MDC-AAF, indicating MT-*Ca*Kip3-MDC-AAF has the most mobile loop-2. In both complexes, much of loop-2 is shifted away from helix-12 of α-tubulin to a degree that many of the bonds observed in the APO and AMP-PNP states would not form (**Fig. 3e, f; right panels**). These changes in the conformation of loop-2 are accompanied by movement of the β1 strand and helix-6 of *Ca*Kip3 away from α-tubulin, opening the space needed for the neck-linker to dock against the motor core (**Fig. 3e, f; left panels**).

### Loop-2 modulates CaKip3’s catalytic function according to tubulin protofilament shape

The fact that microtubule-bound *Ca*Kip3 showed more extensive loop-2-tubulin interactions in the open APO- and ANP-states compared to the closed AAF-state suggests that the interaction of loop-2 with the microtubule lattice restricts rotation of the minus subdomain of the motor domain (**Fig. 2 and Fig. 3**). To investigate this hypothesis experimentally, we designed and expressed a mutant version of the *Ca*Kip3-MDN construct, whose long loop-2 was replaced with the short loop-2 sequence found in human kinesin-1, *Hs*KIF5B (**Supplementary Fig. 1a; *Ca*Kip3-MDN_L2-*Hs*KHC_**), which does not contact α-tubulin in any of the available microtubule-bound or tubulin-bound kinesin-1 structures^14,33,34^. In line with our hypothesis, the cryo-EM structure of this *Ca*Kip3 construct bound to microtubules has a docked neck-linker and closed nucleotide pocket in the ATP-like state (**Fig. 4b; MT-*Ca*Kip3-MDN_L2-*Hs*KHC_-ANP**), unlike the *Ca*Kip3-MDN construct with a full loop-2 (**Fig. 4c; MT-*Ca*Kip3-MDN-ANP**). These data show that the kinesin-8-specific elongated loop-2 acts as a tubulin tether on the microtubule lattice pre-ATP hydrolysis, preventing the motor from closing, and that removing the *Ca*Kip3 loop-2 residues that interact with the microtubule lattice enable the motor domain to form the closed conformation in the ATP-bound state. The MT-*Ca*Kip3-MDN_L2-*Hs*KHC_-ANP structure also suggests that loop-2 of kinesin-8 is directly involved in inducing tubulin curvature because the tubulin conformational changes in this structure are less extensive than in the MT-*Ca*Kip3-MDC-AAF structure (**Fig. 4d, Supplementary Fig. 6d, f; compare blue arrows on α-tubulin**).

To learn how the loop-2-tubulin interactions and motor domain conformations observed for microtubule-bound *Ca*Kip3 relate to its motile activity, we assessed the microtubule gliding velocity of the *Ca*Kip3-MDN and *Ca*Kip3-MDN_L2-*Hs*KHC_ motors. The gliding data shows that elimination of the *Ca*Kip3 loop-2 increased the microtubule-gliding speed of the motor by over three-fold (**Fig. 4e, g, Supplementary Fig. 7a**). Conversely, when loop-2 of *Ca*Kip3 was grafted into an equivalent kinesin-1 construct (*Hs*KHC-MDN_L2-*Ca*Kip3_), the microtubule gliding speed slowed two-fold (**Fig. 4f, g, Supplementary Fig. 7a**). One explanation for these results is that the interaction of loop-2 with straight αβ-tubulin protofilaments delays hydrolysis of ATP by the motor domain, thereby limiting the neck-linker docking event that enables forward stepping. An alternative, but not mutually exclusive explanation, is that the loop-2-tubulin interaction limits the stepping velocity of *Ca*Kip3 by increasing microtubule affinity. To distinguish between these two possibilities, we assessed the microtu-bule-stimulated ATPase activity of *Ca*Kip3-MDN, *Ca*Kip3-MDN_L2-*Hs*KHC_, *Hs*KHC-MDN, and *Hs*KHC-MDN_L2-*Ca*Kip3_ at steady state, and then measured the microtubule affinity of these proteins using a sedimentation assay. The data from the ATPase experiments show that *Ca*Kip3-MDN_L2-*Hs*KHC_ is ~1.6-fold faster at hydrolyzing ATP on microtubules than *Ca*Kip3-MDN (**Fig. 4h, j**). Likewise, the microtubule-stimulated ATP turnover by *Hs*KHC-MDN_L2-*Ca*Kip3_ was almost two-fold slower than *Hs*KHC-MDN (**Fig. 4i, j**). Our microtubule-binding data showed that *Ca*Kip3-MDN has ~35-fold higher affinity for microtubules than *Ca*Kip3-MDN_L2-*Hs*KHC_ when AMP-PNP occupies the nucleotide-binding pocket (**Fig. 4k, m, Supplementary Fig. 7b**). The *Hs*KHC-MDN_L2-*Ca*Kip3_ construct also had a 7-fold greater affinity for the microtubule in the ATP-like state compared to *Hs*KHC-MDN (**Fig. 4l, m, Supplementary Fig. 7c**). These results demonstrate that the extended loop-2 of *Ca*Kip3 both increases the lifetime of the microtubule-bound state and limits ATPase activity of the motor domain on the microtubule lattice.

To understand the functional effects of the loop-2-tubulin interactions observed in the curved tubulin-bound structure of *Ca*Kip3, we compared the dolastatin-stabilized tubulin-ring (D-ring)-stimulated ATPase activities of *Ca*Kip3-MDN and *Ca*Kip3-MDN_L2-*Hs*KHC_. Our data shows that ATP hydrolysis by *Ca*Kip3-MDN_L2-*Hs*KHC_ was two-fold slower than *Ca*Kip3-MDN on D-rings (**Fig. 5a, d**). This reduction in ATPase activity without the full loop-2 region does not appear to be limited by D-ring binding ability because we observed a large proportion of the total *Ca*Kip3-MDN_L2-*Hs*KHC_ protein sedimented with D-rings using a co-sedimentation analysis (**Supplementary Fig. 8a**). Together with our structural data, these results show that the extended *Ca*Kip3 loop-2 functions like a tubulin protofilament shape sensor that modulates the motility and ATPase activity of the motor domain according to the shape of the tubulin protofilament it binds.

**Fig. 5:**
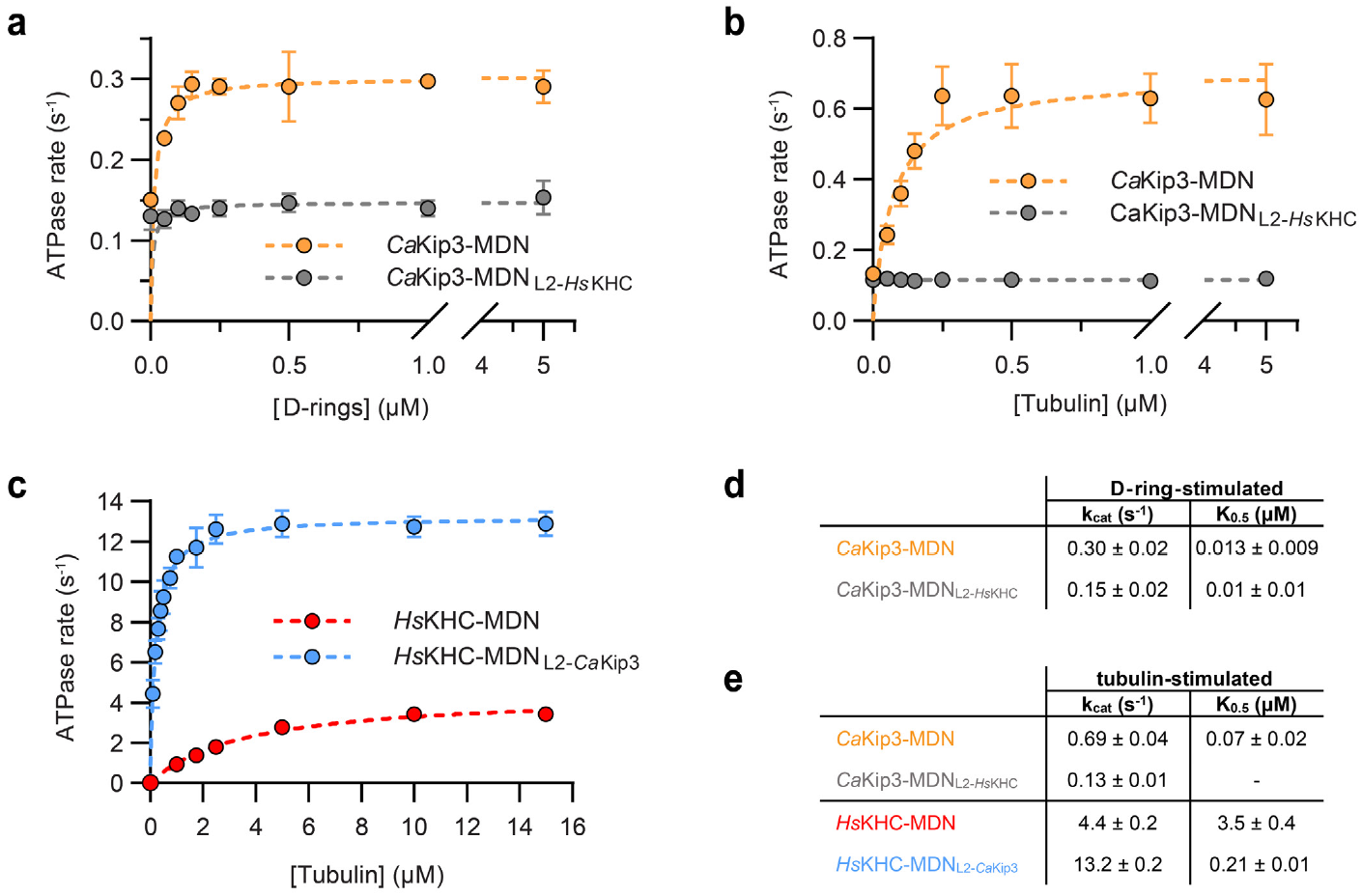
Activity of loop-2 swap mutants on curved tubulin. **(a)** Dolastatin-induced tubulin-ring (D-ring)-stimulated ATPase kinetics of *Ca*Kip3-MDN and *Ca*Kip3-MDN_L2-*Hs*KHC_ at steady state (n=3). **(b)** Free tubulin dimer-stimulated ATPase kinetics of *Ca*Kip3-MDN and *Ca*Kip3-MDN_L2-*Hs*KHC_ at steady state (n=3). **(c)** Free tubulin dimer-stimulated ATPase kinetics of *Hs*KHC-MDN and *Hs*KHC-MDN_L2-*Ca*Kip3_ at steady state (n=3). **(d)** Summary of the D-ring-stimulated ATPase activities of the motors. **(e)** Summary of the tubulin-stimulated ATPase activities of the motors. Values are presented as mean ± SEM. Data were fit to the Michaelis-Menten equation using GraphPad Prism to obtain *K*_0.5, MT_ and *k*_cat_ values.

When we explored the tubulin shape-sensing ability of *Ca*Kip3’s loop-2 further using unpolymerized free tubulin, which is also thought to be in the curved conformation^35,36^, we observed similar experimental results. *Ca*Kip3-MDN_L2-*Hs*KHC_ exhibited almost no free tubulin-stimulated activity, while *Ca*Kip3-MDN had a free tubulin-stimulated ATPase activity that is comparable to other kinesin-8s (**Fig. 5b, e**)^1,24^. Moreover, insertion of the *Ca*Kip3 loop-2 into *Hs*KHC increased free tubulin-stimulated ATPase activity three-fold (**Fig. 5c, e**); opposite its effect on microtubule-stimulated ATPase activity (**Fig. 4i, j**). These changes in tubulin-stimulated ATPase activity, brought about by swapping the loop-2 region, do not appear to be fully explained by a lack of tubulin binding because a large proportion of the total *Ca*Kip3-MDN_L2-*Hs*KHC_ protein eluted with DARPin-capped αβ-tubulin during size-exclusion chromatography (SEC) analysis (**Supplementary Fig. 8b; fractions 13.5 – 15 mL**). However, loop-2-tubulin contacts observed in our *Ca*Kip3 structures do appear to be important tubulin-affinity determinants because insertion of *Ca*Kip3’s loop-2 into kinesin-1 (*Hs*KHC-MDC_L2-*Ca*Kip3_) increased the amount of motor that co-eluted with αβ-tubulin-DARPin relative to *Hs*KHC-MDC (**compare Supplementary Fig. 8c and d; fractions 12 – 12.5 mL**).

### Loop-1 and loop-2 are key drivers of microtubule-depolymerization

In addition to the altered loop-2-tubulin interface and closed conformation of the motor domain of *Ca*Kip3 bound to curved tubulin (CT-*Ca*Kip3-MDC-ANP), a striking finding in this complex is the appearance of map density for loop-1 against the minus end neighboring *Ca*Kip3 motor domain (**Fig. 1f**). This head-to-tail interaction of *Ca*Kip3s is formed by the long polyasparagine repeat region of loop-1 (**Supplementary Fig. 1a**), which inserts into the deep groove between the loop-8 lobe and the core β-sheet of the minus end neighbor motor domain (**Fig. 6a, b**). When the *Ca*Kip3 nucleotide-binding pocket closes upon binding curved tubulin and the inlet and outlet strands of the loop-8 lobe move towards helix-12 of β-tubulin (**Fig. 2d**), this cavity opens, and loop-1 of the plus end neighbor kinesin inserts and interacts with the minus end neighbor kinesin motor domain. Another remarkable feature is the formation of an α-helix in the sequence that precedes the polyasparagine region (**Fig. 6a, right panel**). This helix is an extension of helix-0 and likely forms when the polyasparagine region is stabilized against the adjacent motor. These interactions demonstrate that loop-1 and loop-8 facilitate oligomerization of *Ca*Kip3 upon binding a curved tubulin protofilament. Weaker loop-1 density is also present in the MT-*Ca*Kip3-MDC-AAF complex above the loop-8 lobe and likely represents a similar loop-1-mediated motor-motor interaction (**Supplementary Fig. 9**). No equivalent loop-1 density is observed in the MT-*Ca*Kip3-MDC-APO or -ANP structures. Taken together, these data suggest that loop-1 interactions with a minus end neighbor motor domain are stabilized by *Ca*Kip3 adopting a closed conformation, and that insertion of loop-1 into the other motor domain could potentially stabilize its closed conformation.

**Fig. 6:**
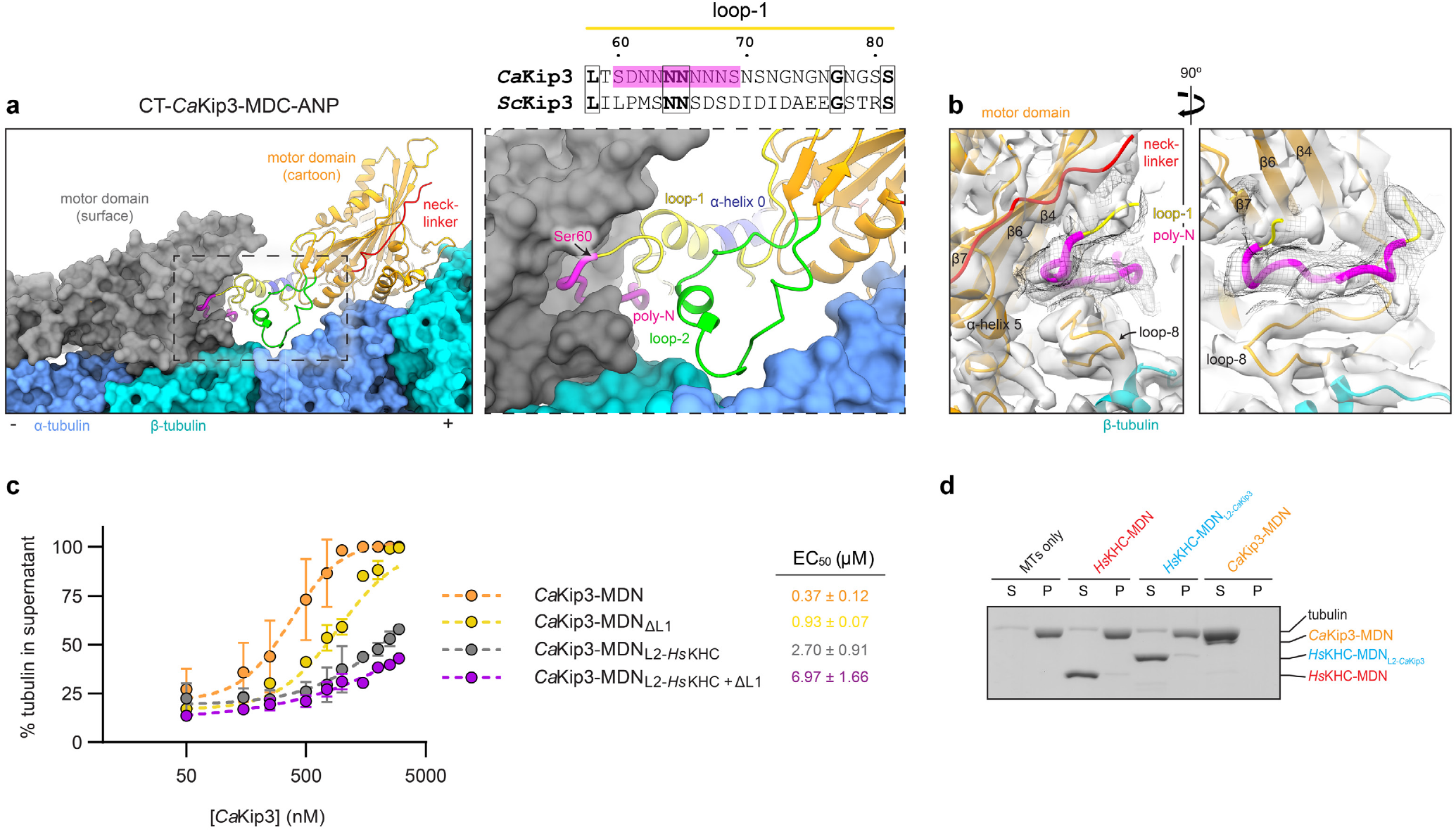
Loop-1 and loop-2 are major contributors to microtubule depolymerization activity. **(a)** Cartoon representation of *Ca*Kip3 in the CT-*Ca*Kip3-MDC-ANP complex showing the polyasparagine section of loop-1 (magenta) inserted into the preceding motor domain (shown in surface representation). The insert shows “close up” view of the polyasparagine section in order to visualize structural elements involved in complex formation. Sequence alignment shows polyasparagine section of loop-1 in *Ca*Kip3 compared to *Sc*Kip3. **(b)** Cryo-EM map of the CT-*Ca*Kip3-MDC-ANP complex. A low-pass filtered map (5 Å) is shown as a black mesh around the polyasparagine track of loop-1. **(c)** Microtubule depolymerization by sedimentation dose-response curves for the indicated *Ca*Kip3 proteins. 2 µM GMP-CPP-stabilized microtubules were incubated with increasing concentrations of *Ca*Kip3-MDN, *Ca*Kip3-MDN_∆L1_, or *Ca*Kip3-MDN_L2-*Hs*KHC_, *Ca*Kip3-MDN_L2-*Hs*KHC+∆L1_ in the presence of ATP. The data from three independent experiments were analysed and fit to the four-parameter logistic equation. EC_50_ values are presented as the mean ± SEM. **(d)** Depolymerization of 2 µM GMP-CPP-stabilized microtubules by 3 µM of *Hs*KHC-MDN, 3 µM *Hs*KHC-MDN_L2-*Ca*Kip3_, and 1 µM *Ca*Kip3-MDN assessed by sedimentation. Reactions were incubated for 20 minutes in the presence of 20 mM MgATP, then free tubulin and microtubule polymers were separated into supernatant (S) and pellet (P) fractions by ultra-centrifugation. Equal portions of (S) and (P) were loaded and analyzed on a 10% Coomassie blue-stained SDS-PAGE gel.

Using a sedimentation assay that separates microtubules from soluble tubulin subunits, we monitored GMP-CPP-stabilized microtubule depolymerization by *Ca*Kip3-MDN, a *Ca*Kip3-MDN construct lacking loop1 (*Ca*Kip3-MDN_ΔL1_), the loop-2 swap construct (*Ca*Kip3-MDN_L2-*Hs*KHC_), and a *Ca*Kip3-MDN_L2-*Hs*KHC_ construct lacking loop-1 (*Ca*Kip3-MDN_L2-*Hs*KHC+ΔL1_). Analysis of these reactions by SDS-PAGE revealed that loop-1 and loop-2 are both important for *Ca*Kip3’s microtubule-depolymerase activity, and that loss of loop-2 causes the most impairment of this activity (**Fig. 6c, Supplementary Fig. 10**). *Ca*Kip3-MDN was 2.5-fold more active than *Ca*Kip3-MDN_ΔL1_ and 7-fold more active than *Ca*Kip3-MDN_L2-*Hs*KHC_ at depolymerizing microtubules (**Fig. 6c**). Loss of loop-1 and loop-2 simultaneously (*Ca*Kip3-MDN_L2-*Hs*KHC+ΔL1_) resulted in a 19-fold decrease in microtubule depolymerization activity (EC_50_). These dramatic effects of loop-1 and loop-2 truncation support the functional importance of these regions in the microtubule depolymerization activity of *Ca*Kip3.

The small amount of microtubule depolymerization activity retained by *Ca*Kip3-MDN_L2-*Hs*KHC+ΔL1_ shows that loop-1 and loop-2 are not the only features of the motor domain involved in catalyzing this process. The inability of *Hs*KHC-MDN_L2-*Ca*Kip3_ to depolymerize microtubules (**Fig. 6d**) and the tubulin curvature that we observe in the *Ca*Kip3-MDN_L2-*Hs*KHC_-ANP complex further support this idea. Overall, our structural data show that the closed nucleotide-binding pocket conformation of the motor domain actively induces tubulin curvature, which likely accounts for the residual depolymerization activity of the *Ca*Kip3-MDN_L2-*Hs*KHC+ΔL1_ construct. Cataloguing the differential tubulin contacts made by the open and closed conformations of the motor highlights that several closed-conformation-specific contacts are formed besides those in loop-2 (**Supplementary Fig. 11**). While many of these interactions likely contribute to the bending of tubulin, loop-2 adds to the tubulin-curving ability of the motor. This is supported by the greater degree of α-tubulin displacement relative to β-tubulin between the MT-*Ca*Kip3-MDC-ANP and MT-*Ca*Kip3-MDC-AAF complexes than between the MT-*Ca*Kip3-MDN_L2-*Hs*KHC_-APO and MT-*Ca*Kip3-MDN_L2-*Hs*KHC_-ANP complexes (**Supplementary Fig. 6d, f**). Moreover, the loss of microtubule depolymerization activity caused by elimination of loop-1 (*Ca*Kip3-MDN_ΔL1_) indicates that loop-1’s interactions with the minus end neighboring motor domain are also involved in the microtubule depolymerization mechanism of *Ca*Kip3. These interactions may allow the linked motor domains to act cooperatively (**Fig. 1f, Supplementary Fig. 3b**).

## Discussion

### Summary of the kinesin-8 mechanism

The central focus of this research was to understand the mechanistic basis for the dual ability of kinesin-8s motors to move processively on microtubules and to depolymerize microtubule plus-ends. To achieve this, we obtained cryo-EM structures that demonstrate how the unique features of a kinesin-8 motor domain interact with αβ-tubulin subunits in the microtubule lattice and on microtubule ends. We also obtained functional data to show how the motor domain conformation and catalytic activity affects, and is affected by, the distinctive shapes of these tubulin polymers. Our findings demonstrate that, when kinesin-8 encounters curved tubulin protofilaments, or protofilaments that can become curved, which are found at the microtubule plus end, its motor domain readily forms a pro-depolymerization state. In this state, the elongated loop-2 region moves toward the motor core to accommodate the position of α-tubulin in curved αβ-tubulin subunits (**Fig. 7**). This displacement of loop-2 is accompanied by closing of the ATP pocket to form a nucleotide-hydrolysis-competent active site and a docked neck-linker. In addition, loop-1 becomes more structurally ordered and inserts into the deep groove between loop-8 and the core β-sheet of the preceding motor domain. By maintaining loop-2 contacts with tubulin, and the loop-1 linkage between motors, the conformational transition of multiple motor domains to a closed conformation could increase tubulin protofilament curvature enough to trigger microtubule depolymerization. Alternatively, when kinesin-8 binds tubulin protofilaments that are constrained to a straight conformation by the lateral protofilament interactions of the microtubule lattice, the interactions between loop-2 and α-tubulin restrict the motor domain from transitioning to the closed conformation with a docked neck-linker. Unable to stably interact with the preceding motor via loop-1, or to curve tubulin enough to disrupt protofilament contacts, some of the loop-2-tubulin bonds eventually break, allowing the motor domain to form the pro-motility state, in which the neck-linker docks and the tethered head is moved to the next αβ-tubulin subunit.

**Fig. 7:**
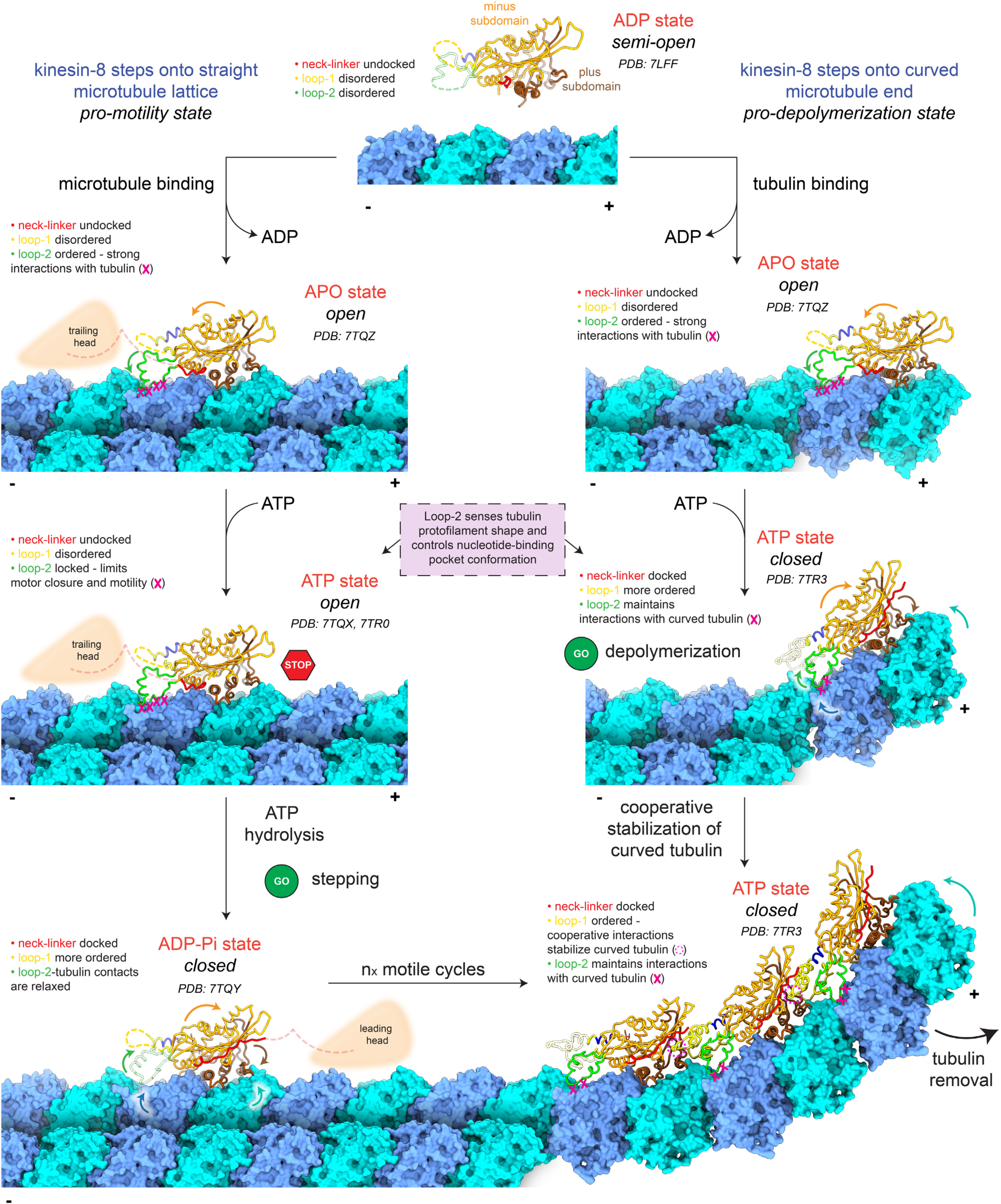
Model for tubulin shape-induced alternations between pro-motility and pro-depolymerization states of kinesin-8.

### The closed conformation induces tubulin curvature

Our structural results demonstrate that the closed conformation of the *Ca*Kip3 nucleotide-binding pocket not only accommodates the curved conformation of tubulin but also actively induces tubulin curvature (**Fig. 2, Fig. 3, Fig. 4, Supplementary Fig. 6**). Comparing the microtubule structures with the *Ca*Kip3 motor domain in the open or closed conformations indicates that motor domain closure is accompanied by tubulin structural changes toward the curved conformation. This conformational change occurs with or without an intact loop-2 but appears more extensive in the latter case. These results indicate that loop-2 is important but not essential for kinesin-8 induced tubulin bending. This could explain the residual depolymerization activity we and others observe with kinesin-8 constructs with a mutated or deleted loop-2. (**Fig. 6c**)^20,24^. Previously, it was reported that loop-11 acts to recognize the curved conformation of tubulin by selectively interacting with an aspartate residue on α-tubulin when tubulin is curved (Asp118 – yeast tubulin, Asp116 – bovine or porcine tubulin)^20^. This interaction was proposed to suppress the ATPase activity of the motor at the microtubule plus-end, leading to prolonged plus-end binding and subsequent microtubule depolymerization. In our CT-*Ca*Kip3-MDC-ANP structure, loop-11 residues are not within bonding distance of Asp116/118 and we observe no structural evidence of this interaction (**Supplementary Fig. 12**). While we cannot rule out the possibility that the conformation of loop-11 on curved tubulin may limit ATPase activity, our structures suggest Asp116/118 is not a major tubulin contact.

### Comparing the mechanochemical cycles of motile motors, depolymerizers, and motile depolymerizers

We observe that the mechanochemical cycle of a motile depolymerase is distinct from that of a strictly motile or strictly depolymerizing kinesin. During motility, the *Ca*Kip3 nucleotide-binding pocket remains in the open conformation following ATP-binding. This is similar to the conformation of the depolymerizing kinesin-13s following ATP binding, and dissimilar from motile motors whose nucleotide pockets close in the equivalent nucleotide state. The fact that kinesin-8s and kine-sin-13s both have elongated loop-2 regions that make microtubule contacts is further support of the hypothesis that loop-2 is also responsible for the open nucleotide-binding pocket of AMP-PNP-bound kinesin-13^16^. On the other hand, a promotility microtubule bound state for kinesin-13s with a closed nucleotide binding pocket and docked neck linker has not been reported. Differences between the loop-2 of kinesin-8s and kinesin-13s appear well suited for the different functionalities of these kinesins. The rigid kinesin-13 loop-2 inserts into the tubulin inter-dimer interface and make similar contacts whether tubulin is in the straight or curved tubulin conformations^16,19^. However, to maintain these contacts the motor domain must be in the open conformation when interacting with straight tubulin and in the closed conformation when interacting with curved tubulin. In the case of kinesin-8s a less structured loop-2 can form alternate contacts with tubulin allowing the motor domain to close and reach the pro-motility state while bound to straight tubulin in the microtubule lattice.

While the loop-2 region of kinesin-13s has a highly conserved motif (KVD) that is critical for microtubule depolymerization^30,32,37^, the loop-2 of kinesin-8s is not highly conserved. As we show here, and as others have previously reported, it is important but not essential for microtubule depolymerization (**Fig. 6c, d**)^20,24^. Our results also agree with previous experiments demonstrating that mutations to the kinesin-8 loop-2 can impart faster motility of monomeric constructs^24^, and that kinesin-1 grafted with enough kinesin-8 parts (loop-2, loop-11, and the neck-linker) has depolymerization activity but is a slower motor^20,24^. We now build on these previous findings and clarify the role of loop-2 as a structural element that controls whether *Ca*Kip3 takes on a pro-motility or pro-depolymerization conformation, based on tubulin curvature.

### Establishing the structural basis for cooperativity

Other studies of *Sc*Kip3 showed that kinesin-8 acts cooperatively to mediate length-dependent microtubule depolymerization^2,4^. Our structure of *Ca*Kip3 bound to D-rings provides a possible explanation for the cooperative mechanism of kinesin-8 microtubule depolymerization activity. We show that loop-1 (and loop-2) enable formation of linear arrays of motor domains on curved tubulin protofilaments. α-helix-0 extends toward the ends of the docked neck-linker of the proximal motor domain and then transitions into a turn near Asp55 of loop-1 that inserts into the space between the loop-8 lobe and the core β-sheet. Three segments of loop-1 contact the adjacent motor domain, burying ~1075 Å² of the adjacent motor domain surface. The polyasparagine track in loop-1 (including residues Asn62, Asn64, Asn65, Asn66, Ser69, Asn70, Ser71, Asn78, Gly79) forms the key part of this interaction, and some of this track is conserved in *Sc*Kip3. In *Sc*Kip3, two Asn residues are followed by a Ser/Asp and Ile/Asp repeat in the homologous region (**Fig. 6a, Supplementary Fig. 1a**). Previous studies reported that polyasparagine tracks in loop-8 of yeast kinesin-5, Cin8, may be involved in oligomerization due to the aggregation-inducing effect these sequences can have^38^. The binding site for the polyasparagine track of loop-1 is formed when the loop-8 lobe separates from the core β-sheet of the adjacent motor domain. Lys261, Arg269, Phe326, Phe392, and Arg435 are repositioned and accommodate loop-1. Interestingly, the loop-8 lobe also separates from the motor core in the MT-*Ca*Kip3-MDC-AAF structure, and weak density for loop-1 insertion is visible, indicating that cooperative kinesin-8-mediated tubulin bending can begin on microtubule proto-filaments that have minimal curvature.

In summary, these studies provide a new understanding of how kinesin-8s function as dual-activity motor proteins. Our findings implicate loop-2 as a tubulin protofilament shape sensor that regulates conformational changes in the kinesin-8 motor domain, which can either induce curvature of protofilaments at the microtubule end, leading to microtubule depolymerization, or enable forward displacement of the neck linker for processive motility if the motor domain is bound to protofilaments in the microtubule lattice. They also provide a structural basis for the cooperative depolymerization activity that has been reported for members of this kinesin family.

## Materials and Methods

### Cloning

The genes for *Candida albicans* Kip3 and *Homo sapiens* kinesin heavy chain (*Hs*KHC/Kif5B) were codon-optimized for expression in *E. coli* and ordered as chemically synthesized DNA fragments from Integrated DNA Technologies (IDT). For all kinesin constructs used in these studies (**Supplementary Fig. 1a**), gene fragments of the desired length were PCR amplified and cloned into pET-24d(+) using NcoI and XhoI, yielding a C-terminal 6X-His tag. Mutant genes were generated either through overlap-extension PCR (*Ca*Kip3-MDN_L2-*Hs*KHC_, *Ca*Kip3-MDN_ΔL1_, *Ca*Kip3-MDN_L2-*Hs*KHC+ΔL1_) or ordered as chemically synthesized DNA (*Hs*KHC-MDC_L2-*Ca*Kip3_). The *Ca*Kip3-MDN_L2-*Hs*KHC_ and *Ca*Kip3-MDN_L2-*Hs*KHC+ΔL1_ constructs have the *Hs*KIF5B loop-2 sequence ‘IASK’ (residues I41-K44) replacing *Ca*Kip3 residues F116-H144. The *Ca*Kip3-MDN_ΔL1_ and *Ca*Kip3-MDN_L2-*Hs*KHC+ΔL1_ constructs have residues V44-G101 removed. The *Hs*KHC-MDN_L2-*Ca*Kip3_ and *Hs*KHC-MDC_L2-*Ca*Kip3_ constructs have the *Ca*Kip3 loop-2 sequence from F116-H144 replacing the *Hs*KIF5B loop-2 sequence ‘IASK’ (residues I41-K44). Constructs *Hs*KHC-MDC and *Hs*KHC-MDC_L2-*Ca*Kip3_ contained a cleavable C-terminal TEV-SNAP-6X-His tag (SNAP-tag from NEB). DARPin-D2 was ordered as a codon-optimized gene product from IDT and cloned into pET16b using Gibson Assembly, producing an N-terminal 6X-His tag. All plasmids were sequenced to verify correct assembly.

### Expression and purification of recombinant proteins

For all kinesin proteins and DARPin D2, the same general purification scheme was used. Plasmids were transformed into *E. coli* BL21(DE3) cells (or BL21 RIL cells for DARPin-D2) and proteins were expressed in Luria-Bertani (LB) media supplemented with the appropriate antibiotic. Protein production was induced with 1 mM isopropyl β-D-1-thiogalactopyranoside (IPTG) and cells were grown overnight between 16-25 °C. Cell pellets were lysed in lysis buffer (10 mM sodium phosphate, 300 mM NaCl, 2 mM MgCl_2_, 1 mM EGTA, 0.2 mM ATP, 5 mM 2-Mercaptoethanol (2-ME), 0.2 mg/mL lysozyme (Bioshop), Pierce Protease Inhibitor Tablets (Thermo), pH 7.2) using sonication. Clarified supernatant was loaded onto Ni-NTA resin (Qiagen) equilibrated in wash buffer (10 mM sodium phosphate, 300 mM NaCl, 2 mM MgCl_2_, 0.1 mM EGTA, 20 mM imidazole, 0.2 mM ATP, 5 mM 2-ME, pH 7.2) and resin was thoroughly washed with wash buffer. Target protein was eluted with elution buffer (wash buffer + 300 mM imidazole) then dialyzed overnight into HEPES buffer (20 mM HEPES, 1 mM MgCl_2_, 150 mM NaCl, 0.2 mM ATP, 1 mM dithiothreitol (DTT), pH 7.2). The next morning, samples were run over a Superdex 200 26/60 size-exclusion column (GE Healthcare) equilibrated with HEPES buffer. Fractions containing target protein were pooled, concentrated, flash frozen in liquid N_2_, and stored at −80 °C.

Tubulin was purified from bovine brains by two cycles of polymerization-depolymerization in a high-molarity PIPES buffer as described by Castoldi *et al*.^39^. Aliquots were flash frozen in liquid N_2_ and stored at −80 °C until use.

### Preparation of taxol-stabilized microtubules and D-rings for biochemical assays

Taxol-stabilized microtubules were assembled as previously described with some modifications^40^. Soluble tubulin was diluted to 5 mg/mL in BRB80 buffer (80 mM PIPES, 1 mM MgCl_2_, 1 mM K-EGTA, 1 mM DTT, pH 6.8) supplemented with 2 mM GTP and depolymerized on ice for 10 minutes. The mixture was clarified by centrifugation at 90,000 rpm (TLA-100; Beckman Coulter) for 5 minutes at 4 °C, then microtu-bule-polymerization was induced by addition of 10% DMSO and incubation at 37 °C. After 40 minutes, 20 μM taxol (paclitaxel-d5, Toronto Research Chemicals) was added to the reaction mixture and incubation was continued at 37 °C for another 40 minutes. Microtubules were sedimented at 60,000 rpm (TLA-100; Beckman Coulter) for 15 minutes at 25 °C and resuspended in the appropriate reaction buffer supplemented with 20 μM taxol. Unless otherwise specified, microtubule concentration was determined by measuring optical density at 280 nm (A_280_). To mitigate the light scattering effect of microtubule polymers, microtubules were depolymerized by incubation on ice and addition of 5 mM CaCl_2_ prior to measuring A_280_.

Dolastatin-induced tubulin-rings (D-rings) were polymerized by diluting tubulin to 40 μM in BRB80 buffer containing 80 μM dolastatin-10 (APExBIO) and incubating the reaction for 1 hour at 25 °C. Whenever D-rings were used in experiments, the concentration of dolastatin present was at least two-fold more than the concentration of tubulin and never below a concentration of 20 μM. Assembled D-rings were used within 2 hours.

### ATPase assay

ATP turnover by kinesins was measured when stimulated by taxol-stabilized microtubules, D-rings, or soluble tubulin. To prepare tubulin for the ATPase assays, tubulin stock was diluted to 1.7 mg/mL in A25 buffer (25 mM ACES, 2 mM MgAcetate, 2 mM K-EGTA, 0.1 mM K-EDTA, 1 mM 2-ME, pH 6.9), incubated on ice for 10 min, and clarified at 90,000 rpm (TLA-100; Beckman Coulter) for 5 min at 4 °C. ATP turnover was measured using an enzyme-coupled system^41,42^ and in specific cases the EnzChek Phosphate assay (Thermo Fisher) was also used to verify results (for tubulin and D-ring-stimulated *Ca*Kip3-MDN and *Ca*Kip3-MDN_L2-*Hs*KHC_). For the enzyme-coupled system, reactions were prepared in A25 buffer containing 1 mM MgATP, 2 mM phosphoenol-pyruvate (K-PEP), 0.25 mM NADH, 60 μg/mL lactate dehydrogenase, 60 μg/mL pyruvate kinase, 20 μM taxol (for microtubule reactions), 20 μM dolastatin (for D-ring reactions), microtubules or D-rings or tubulin, and 200 nM kinesin (for kinesin-8 reactions) or 15 nM kinesin (for kinesin-1 reactions). Reactions containing soluble tubulin also included 0.5 mM GDP to hinder microtubule-polymerization. For the EnzChek Phosphate assay, reactions were prepared in A25 buffer and contained 2 mM MgATP, 200 μM 2-amino-6-mercapto-7-methylpurine riboside (MESG), 1 unit/mL purine nucleoside phosphorylase (PNP), 20 μM dolastatin (for D-ring reactions), D-rings or tubulin, and 200 nM kinesin-8. Using a SpectraMax iD3 plate reader (Molecular Devices), A_340_ values (enzyme-coupled system) or A_360_ values (Enzchek phosphate assay) were collected in 15-second intervals for 15 minutes. Rates of ATP turnover were determined from the initial linear portion of the data and fit to the Michaelis-Menten equation using GraphPad Prism 8.0 (GraphPad Software, San Diego, CA) to determine K_0.5_ and *k*_cat_.

### Microtubule co-sedimentation assay

Microtubule-kinesin co-sedimentation reactions were prepared and analyzed as described^41^ with the following modifications. Prior to setting up co-sedimentation reactions, kinesin motor was pre-cleared by centrifugation at 90,000 rpm (TLA-100; Beckman Coulter) for 5 minutes at 4 °C. Co-sedimentation reactions were prepared in 100 μL volumes of BRB80 buffer supplemented with 100 mM KCl and contained 2 mM adenylyl-imidodiphosphate (AMP-PNP), 1 μM kinesin, 20 μM taxol, and 0-10 μM microtubules. Reaction mixtures were incubated for 20 minutes at 25 °C then centrifuged at 60,000 rpm (TLA-100; Beckman Coulter) for 15 minutes at 25 °C. The 50 μL of the supernatants were retained, and the remainder was discarded. Pellets were resuspended in 100 μL BRB80 buffer and 50 μL was retained for SDS-PAGE analysis. Supernatant and pellet samples (10 µL per lane) were loaded on 10% SDS-PAGE gel, ran, and then were visualized by staining with Coomassie brilliant blue R-250. Band intensities were quantified with ImageJ and the percentage of kinesin complexed with the microtubule was plotted as a function of microtubule concentration. To obtain values for K_d_, data was fit to a quadratic equation using GraphPad Prism 8.0: *(MT · E)/E = 0.5(E_0_ + K_d,MT_ + MT_0_) − [(E_0_ + K_d,MT_ + MT_0_)^2^ − (4E_0_MT_0_)^1/2^]*, where *E_0_* is the total kinesin concentration, *MT·E* is the amount of kinesin bound to the microtubules, *MT_0_* is the total microtubule concentration, and *K*_d_ is the dissociation constant.

### D-ring co-sedimentation assay

D-ring co-sedimentation assays were performed as described^43^ with the following modifications. Kinesin motor was pre-cleared by centrifugation at 90,000 rpm (TLA-100; Beckman Coulter) for 5 minutes at 4 °C. Co-sedimentation reactions were prepared in 100 μL volumes of BRB80 buffer contained 4 μM D-rings, 20 μM Dolastatin, 4 μM kinesin, and 2 mM AMP-PNP. Dolastatin was also present in control reactions that did not contain D-rings. Reactions were incubated for 10 minutes at 25 °C, then centrifuged at 100,000 rpm (TLA-100; Beckman Coulter) at 25 °C for 15 minutes. The top 50 μL of the reaction supernatants were retained and the remainder was discarded. Pellets were resuspended in 100 μL BRB80 buffer and 50 μL was retained for SDS-PAGE analysis. Supernatant and pellet samples were analyzed on SDS-PAGE gels stained with Coomassie brilliant blue R-250.

### Motility assay

Rhodamine-labelled taxol-stabilized microtubules were assembled as previously described with the following changes^44^. Rhodamine-labelled porcine tubulin (Cytoskeleton Inc.) and unlabelled tubulin were mixed in a 1:7 molar ratio (5 mg/mL total tubulin) in BRB80 buffer supplemented with 10% DMSO and 2 mM GTP. The mixture was depolymerized on ice for 10 minutes, then clarified at 90,000 rpm (TLA-100; Beckman Coulter) for 5 minutes at 4 °C. The supernatant was polymerized at 37 °C for 30 minutes, then diluted two-fold with BRB80 buffer containing 40 μM taxol. The incubation was continued for another 30 minutes at 37 °C, then microtubules were sedimented at 60,000 rpm at 25 °C for 15 minutes. The microtubules contained in the pellet were carefully resuspended in BRB80 buffer containing 40 μM taxol.

Flow chambers for the motility assay were constructed by adhering siliconized coverslips (Hampton Research) to microscopy slides using double-sided tape. The total chamber volume was roughly 8 μL. Three oxygen-scavenging mixtures (OSM) were prepared fresh for each experiment: OSM-0 (BRB80, 1.5 mM MgAc, 2 mM 2-ME, 200 μg/mL glucose oxidase, 175 μg/mL catalase, 25 mM glucose), OSM-1 (OSM-0 with addition of 1.5 mM AMP-PNP, 40 μM taxol), and OSM-2 (OSM-0 with 1.5 mM MgATP, 40 μM taxol). Stock anti-His antibody (Thermo Fisher) was diluted to 70 μg/mL in BRB80 buffer, flowed through the chamber, and allowed to adhere for 5 minutes. Chambers were washed with BRB80 buffer, then 1% F-127 pluronic acid in BRB80 buffer was added to block non-specific binding of microtubules. After a 10-minute incubation, chambers were washed with BRB80 buffer and 0.5 μM His-tagged kinesin diluted in OSM-0 was introduced into the chamber. Kinesin was allowed to bind to anti-His antibodies for 10 minutes, then a 1:60 dilution of rhodamine-labelled microtubules in OSM-1 was flowed into the chamber. After a 5-minute incubation, OSM-2 was flowed into the chamber to remove unbound microtubules and initiate the reaction. Imaging was performed at 25 °C on an Olympus IX83 inverted fluorescence microscope equipped with an Andor Zyla 4.2 Plus camera. Images were collected every 20 seconds for kinesin-8 timelapses and every 5 seconds for kinesin-1 timelapses. Microtubule velocity was calculated using ImageJ (NIH) by tracking the position of the microtubule end in each frame. The average distance traveled between each frame was divided by the time increment between frames to yield a velocity value for each microtubule. Histograms fit with Gaussian curves were generated in GraphPad Prism 8.0.

### Microtubule-depolymerization assay

Assessment of microtubule-depolymerization by sedimentation was carried out as described with minor changes^24^. To assemble microtubules stabilized with guanylyl-(a,(3)-methylene-diphosphonate (GMP-CPP) (Jena Bioscience), tubulin was diluted to 2 mg/mL in BRB80 buffer supplemented with 0.5 mM GMP-CPP. The mixture was incubated on ice for 10 minutes then clarified at 90,000 rpm (TLA-100; Beckman Coulter), 5 minutes, 4 °C. The supernatant was polymerized at 37 °C for 1 hour, then microtubules were sedimented at 60,000 rpm (TLA-100; Beckman Coulter) for 15 minutes at 25 °C. Resuspended microtubules were then depolymerized, clarified, and re-polymerized to increase the occupancy of GMP-CPP within the microtubule lattice. The second microtubule-polymerization step was performed with 1 mM GMP-CPP. The final GMP-CPP-stabilized microtubules were sedimented at 60,000 rpm (TLA-100; Beckman Coulter) for 15 minutes at 25 °C and resuspended in BRB80 buffer.

Sedimentation reactions were prepared as 50 μL of BRB80 buffer containing 2 μM GMP-CPP-stabilized microtubules, 20 mM MgATP and 0-3 μM kinesin. Reactions were incubated for 20 minutes at 25 °C then centrifuged at 60, 000 rpm (TLA-100; Beckman Coulter) for 15 minutes, 25 °C. The top 30 μL of the reaction supernatant was retained and the remainder was discarded. Pellets were resuspended in 50 μL BRB80 buffer and 30 μL was retained for SDS-PAGE analysis. Supernatant and pellet fractions were visualized by SDS-PAGE stained with Coomassie brilliant blue R-250. The intensity of tubulin in each fraction was quantified with ImageJ (NIH). The percentage of tubulin in the supernatant fraction was plotted as a function of kinesin concentration and values were fit to a four-parameter logistic equation using GraphPad Prism 8.0 to determine relative EC_50_ values.

### Analytical size-exclusion chromatography

Analytical SEC experiments were performed as described previously^19^ with minor changes. A 200 μL reaction was prepared in HEPES buffer (20 mM HEPES, 1 mM MgCl_2_, 150 mM NaCl, 1 mM DTT, pH 7.2) containing 23 μM kinesin, 23 μM tubulin, 23 μM DARPin-D2, 0.1 mM GDP, and 0.2 mM AMPPNP. Reactions were incubated on ice for 30 minutes, clarified at 13,148 × g for 10 minutes at 4 °C, then 100 μL of the mixture was loaded onto a Superdex 200 10/300 GL size-exclusion column (GE Healthcare) equilibrated in HEPES buffer. Fractions of interest (0.5 mL each) were concentrated using an Amicon Ultra filter with a 10 kDa cutoff for 10 min at 13,148 × g, 4 °C and the resulting volume was made up to 60 μL. Fractions were visualized by SDS-PAGE stained with Coomassie brilliant blue R-250. For specific samples, immunoblotting using HRP-conjugated anti-His antibody (Abcam, ab1187) was performed based on an established protocol^45^. Following SDS-PAGE of fractions diluted 1:20, a wet electroblot at 400 mA for 2 hours was used to transfer proteins to a PVDF membrane. Membranes were blocked with 5% milk powder in TBST buffer (50 mM Tris, 150 mM NaCl, 0.05% Tween 20, pH 7.5) for 1 hour. Membranes were incubated in anti-His antibody (diluted 1:1000 in blocking buffer) for 1 hour. Membranes were washed 3 times with TBST at room temperature then incubated with the Pierce Enhanced Chemiluminescence Western Blotting Substrate (Thermo Fisher) prior to imaging. Blots were imaged using an Azure C300 Digital Imager.

### Crystallization of CaKip3-MDN-ADP

Crystals of *Ca*Kip3-MDN-ADP were grown using hanging drop vapour diffusion by mixing the protein in a 1:1 ratio with a precipitant solution of 0.1 M MMT (Malic acid, MES and TRIS), 25% PEG 1500, pH 8 at 277 K. The total drop volume was 4 μL. Prior to mixing, the solution of purified kinesin protein (roughly 22 mg/mL) was supplemented with 1 mM ATP. Crystals took approximately one week to grow full-sized. Before collecting diffraction data, crystals were transferred to a cryoprotectant solution composed of 0.1 M MMT, 25% PEG 1500, 20% ethylene glycol, pH 8, and were flash frozen in liquid N_2_.

X-ray diffraction data for the *Ca*Kip3-MDN-ADP crystals were collected using the synchrotron beamline CMCF 08ID-1 of the Canadian Light Source (Saskatoon, Canada) at 100K. The in-house software program AutoProcess developed at the CMCF was used to run XDS^46^, which processed and scaled the diffraction data and produced the reflection output file for structure determination. The initial *Ca*Kip3-MDN-ADP structure was solved by molecular replacement using the KIF18A motor domain (PDB: 3LRE) as a search model in MOLREP^47^. The final model was produced after several cycles of manual building in COOT^48^ and structure refinement using PHENIX^49^. Diffraction data collection and refinement statistics are summarized in **Table 1**.

### Cryo-EM samples preparation

The datasets were collected on gold grids (UltrAuFoil R2/2 200 mesh) plasma cleaned just before use (Gatan Solarus plasma cleaner, at 15 W for 6 s in a 75% argon/25% oxygen atmosphere).

Fifteen-protofilament-enriched microtubules were prepared from porcine brain tubulin (Cytoskeleton, Inc. CO)^16^. Microtubules were polymerized fresh the day of the cryo-EM sample preparation. Kinesin aliquots were thawed on ice just before use. All nucleotides stock solutions used were prepared with an equimolar equivalent of MgCl_2_ to ADP or AMP-PNP. Four microliters of a microtubule solution at 2–5 μM tubulin in BRB80 buffer supplemented with 20 μM paclitaxel were layered onto the EM grid and incubated for 1 minute at room temperature. This microtubule solution also contains AMP-PNP at 4 mM or ADP at 4 mM plus 2 mM AlCl_3_ and 10 mM KF for the AMP-PNP and ADP-AlF_x_ conditions respectively. During the incubation time, a fraction of the thawed kinesin aliquot was diluted to prepare a 20-μM kinesin solution containing 20 μM paclitaxel and either of the three nucleotide conditions to be probed: (1) 4 mM AMP-PNP, (2) 4 mM ADP plus 2 mM AlCl_3_, and 10 mM KF (ADP-AlF_x_ conditions) or (3) apyrase: 5 × 10^−3^ units per µL (APO conditions). Then the excess microtubule solution was blotted from the grid using a Whatman #1 paper. Four microliters of the kinesin solution were then applied onto the EM grid, transferred to the chamber of a Vitrobot apparatus (FEI-ThermoFisher MA) at 100% humidity where it was incubated for 1 min at room temperature, and blotted for 2.5–3 s with a Whatman #1 paper and −2-mm offset before plunge-freezing into liquid ethane. Grids were clipped and stored in liquid nitrogen until imaging in a cryo-electron microscope.

Dolastatin-10 rings (D-rings) were made by incubating 80 µM porcine brain tubulin (Cytoskeleton, Inc. CO) with 160 µM dolastatin-10 (APExBIO) in BRB80 buffer pH 6.8 and incubating for 1 h at 25 °C. That solution was then diluted to make the cryo-EM sample: 3 µM tubulin D-rings, 20 µM dolastatin-10, 12 µM *Ca*Kip3-MDC, 4 mM AMP-PNP in BRB80 buffer, pH 6.8.

### Cryo-EM data collection

Data were collected at 300 kV on Titan Krios microscopes (**Supplementary Table 1**) equipped with K2 summit detectors for the microtubule datasets and with K3 detector for the CT-*Ca*Kip3-MDC-ANP dataset. Acquisition was controlled using Leginon^50^ with the image-shift protocol and partial correction for coma induced by beam tilt^51^. The defocus ranges and cumulated dose for all datasets are given in **Table 2**.

For the decorated microtubule samples, data collection was performed semi-automatically using one stage shift per hole and 4–6 exposures using image-shift per 2-μm diameter holes. The exposures were fractionated on 40–50 movie frames.

For the CT-*Ca*Kip3-MDC-ANP sample, data collection was performed semi-automatically targeting 5 holes per stage shift and 2 exposures using image-shift per 2-μm diameter holes. The stage was tilted 40 degrees to compensate for preferential orientation of the particles^52^ (Supplementary Fig.3). The other dataset used to make 2D class averages of the structures was collected similarly but with no stage tilt, targeting 5-6 holes per stage shift with 3 exposures per 2-μm diameter holes. The exposures were fractionated on 50 movie frames.

### Processing of the cryo-EM datasets of microtubule-kinesin complexes

The processing was done as previously described^13^. Movie frames were aligned with motioncor2^53^ generating both dose-weighted and non-dose-weighted sums and correcting for magnification anisotropy (**Supplementary Table 1**). Before each of the corresponding cryo-EM session, a series of ~20 micrographs on a cross-grating calibration grid with gold crystals was used to estimate the current magnification anisotropy of the microscope using the program mag_distortion_estimate v1.0^54^. Magnification anisotropy correction was performed within motioncor2 using the obtained distortion estimates (**Supplementary Table 1**). Contrast transfer function (CTF) parameters per micrographs were estimated with Gctf^55^ on aligned and non-dose-weighted movie averages.

Helical reconstruction on 15R microtubules was performed using a helical-single-particle 3D analysis workflow in Frealign^56^, as described previously^13,16^, with each half filament contributing to a distinct half dataset. The box size used are indicated in **Supplementary Table 1**. Per-particle CTF refinement was performed with FrealignX^57^.

To select for tubulins bound to kinesin motors and to improve the resolution of the kinesin-tubulin complexes the procedure HSARC^13^ was used for these one-head-bound states. The procedure follows these steps:

1. Relion helical refinement. The two independent Frealign helical refined half datasets were subjected to a single helical refinement in Relion 3.1^58^ where each dataset was assigned to a distinct half-set and using as priors the Euler angle values determined in the helical-single-particle 3D reconstruction (initial resolution: 8 Å, sigma on Euler angles sigma_ang: 1.5, no helical parameter search).
2. Asymmetric refinement with partial signal subtraction. An atomic model of a kinesin-tubulin complex was used to generate two soft masks using EMAN pdb2mrc and relion_mask_create (low-pass filtration: 30 Å, initial threshold: 0.05, extension: 14 pixels, soft edge: 6 or 8 pixels). One mask (mask_full_) was generated from a kinesin model bound to one tubulin dimer and two longitudinally flanking tubulin subunits while the other mask (mask_kinesin_) was generated with only the kinesin coordinates. The helical dataset alignment file was symmetry expanded using the 15R microtubule symmetry of the dataset. Partial signal subtraction was then performed using mask_full_ to retain the signal within that mask. During this procedure, images were re-centered on the projections of 3D coordinates of the center of mass of mask_full_ (C_M_) using a box size as indicated in **Supplementary Table 1**. The partially signal subtracted dataset was then used in a Relion 3D refinement procedure using as priors the Euler angle values determined form the Relion helical refinement and the symmetry expansion procedure (initial resolution: 8 Å, sigma_ang: 2, offset range corresponding to 3.5 Å, healpix_order and auto_local_healpix_order set to 5). The CTF of each particle was corrected to account for their different position along the optical axis.
3. 3D classification of the kinesin signal. A second partial signal subtraction procedure identical to the first one but using mask_kinesin_ and with particles re-centered on the projections of C_M_ was performed to subtract all but the kinesin signal. The images obtained were resampled to 3.5 Å/pixel and the 3D refinement from step 2 was used to update the Euler angles and shifts of all particles. A 3D focused classification without images alignment and using a mask for the kinesin generated like mask_kinesin_ was then performed on the resampled dataset (8 classes, tau2_fudge: 4, padding: 2, iterations: 175). All datasets but MT-*Ca*Kip3-MDN-ANP and MT-*Ca*Kip3-MDC-AAF classifications produced one major class with a class average where the kinesin motor densities were well-resolved while the others had absent or poorly-resolved kinesin densities. In all cases, only a single motor main domain conformation (open *vs* closed) was found per dataset. In case of the two largest datasets (MT-*Ca*Kip3-MDN-ANP and MT-*Ca*Kip3-MDC-AAF), 2 classes averages had well resolved densities and the particles belonging to these two classes were merged to generate the average structure. These best classes were selected to generate the average kinesin-tubulin structure in the following step.
4. 3D reconstructions with original images (not signal subtracted). To avoid potential artifacts introduced by the signal subtraction procedure, final 3D reconstructions were obtained using relion_reconstruct on the original image-particles without signal subtraction. To obtain a final locally filtered and locally sharpened map, post-processing of the pair of unfiltered and unsharpened half maps was performed as follows. One of the two unfiltered half-map was low-pass-filtered to 15 Å and the minimal threshold value that does not show noise around the microtubule fragment was used to generate a mask with relion_mask_create (low-pass filtration: 15 Å, extension: 10 pixels, soft edge: 10 pixels). This soft mask was used in blocres^59^ on 12-pixel size boxes to obtain initial local resolution estimates. The merged map was locally filtered by blocfilt^59^ using blocres local resolution estimates and then used for local low-pass filtration and local sharpening in localdeblur^60^ with resolution search up to 25 Å. The localdeblur program converged to a filtration appropriate for the tubulin part of the map but over-sharpened for the kinesin part. The maps at every localdeblur cycle were saved and the maps with better filtration for the kinesin part area, with the aim of resolving better the kinesin loops, were selected.

### Processing of the cryo-EM CT-CaKip3-MDC-ANP dataset

The frames were aligned with relion implementation of the motioncor2 algorithm. Contrast transfer function (CTF) parameters per micrographs were estimated with Gctf^55^ on aligned and non-dose-weighted movie averages. The rest of the processing was done using Relion 3.1^58^ unless indicated otherwise.

A first dataset collected with no stage tilt was analyzed with 2D classifications after picking ring-like structures to observe the different structures present. Representative class averages of the different structures that were detected are presented in **Supplementary Fig. 3**. The samples of *Ca*Kip3 mixed with tubulin, dolastatin-10 and AMP-PNP contained both rings of various diameters and spring/spiral-like flexible structures (**Supplementary Fig. 3a, b**). Because the dolastatin-10 rings have a strong preferential orientation (**Supplementary Fig. 3b**), the processing to obtain a 3D structure was done on a dataset collected at a 40-degree stage tilt. The flexible spring/spirals-like structures were avoided during the particle picking to focus on the rings. A subset of ~2k rings with a wide range of out of plane tilt were first picked manually. On data resampled to ~ 4 Å /pix, a series of 3 cycles of 2D classification followed by re-centering on the ring centers was applied to generate the class averages subsequently used as template for automated picking with a low pass filtration of 50 Å. Using the resulting automatically picked particles, 3 cycles of 2D classification, re-centering, and particle extraction were performed to select rings. The predominant rings type which has 14 asymmetric units (called 'C14 rings' from here) was selected. Representative 2D class averages obtained at the end of the third 2D classification are provided in **Supplementary Fig. 3b**. Local CTF parameters estimation for each of these C14 rings was performed using Gctf. These particles were resampled at 1.5 Å/pixel with 512-pixel box size particle-images and then used in a 3D refinement with C1 symmetry (initial resolution 60 Å) using an initial model generated by ab-initio modelling, with its handedness corrected. A 3D classification on data resampled at ~ 3.5 Å/pixel (6 classes, tau2_fudge 2, 25 iterations) was then used to select the rings contributing to the main class characterized by the most regular ring structure (71% of the particles). A 3D refinement was then performed with a C1 symmetry, the initial model mentioned above low pass filtered at 30 Å and without mask. Duplicated particles were then removed, leading to ~38 k C14 rings (**Table 2**). No further particle classification was used from that stage of the processing, with a homogeneous set of C14 rings. A 3D refinement with C14 symmetry was then performed with an initial resolution of 20 Å, a soft mask around the ring (from the previous map with relion_mask_create: low-pass filtration 30 Å, thresold 0.01, extension 25 pixels, soft edge 10 pixels), the solvent_correct_fsc option and local constrains (offset_range 2, offset_step 1, sigma_ang 2) and led a 6.2 Å map of the full ring structure. A CTF refinement step using only brute-force defocus search over a scan range of 3000 Å was performed followed by a 3D refinement with identical parameters as the previous one which improved the resolution to 5.1 Å. A step of particle polishing (with box size of 700 pixels and scaled down to 512 pixels, 1.5 Å/pixel) without training and followed by a 3D refinement (initial resolution 10 Å, a soft mask around the ring made as before, with the solvent_correct_fsc option and local constrains offset_range 2, offset_step 1 & sigma_ang 2) enabled to improve the resolution to 4.8 Å. Repeating the previous polishing and 3D refinements steps led to a 4.7 Å map at 1.5 Å/pixel. These 2 cycles of particles polishing followed by 3D refinements were repeated without binning the data (box size 700 pixels, 1.083 Å/pixel), this time with local constrains offset_range 6 and sigma_ang 5 and led to a full scale C14 ring map at 4.5 Å. A third cycle didn't improve the resolution and led to the final full C14 ring map (**Fig. 1a**).

The processing was continued by doing local alignments focused on fragments of the C14 rings as described in the following. First, to continue the processing locally without losing the benefit from the particle polishing done on the full rings, the polished particles from the previous polishing step were regenerated on larger 900-pixel boxes. Symmetry expansion of the last alignment data based on the C14 symmetry was followed by a relion particle extraction step using the 900 pixels size polished particles-images as input and with re-centering on the center of a mask covering a single asymmetric unit. This enabled to extract from each of these C14 rings in 900-pixel boxes, 14 particles each in 416-pixel boxes and each centered on a different asymmetric unit, for a total of ~531k particles (**Table 2**). The defocus of these particles was updated, considering their different position along the optical axis. A preliminary PDB was used to create with EMAN pdb2mrc and relion_mask create a soft mask covering 3 asymmetric units (low pass filtration 30 Å, initial threshold 0.3, extension 30 pixels, soft edge 6 Å). That mask was used in a focused refinement with fixed priors on the Euler angles and shifts based on the previous alignment data, local constrains (offset_range 6, offset_step 1, sigma_ang 5) and started with the reconstruction of this expanded data filtered at 10 Å. This focused refinement produced a 4.0 Å map (**Fig. 1f, top structure**). Repeating the same focused refinement but with a mask made similarly covering only the central asymmetric unit (low pass filtration 30 Å, initial threshold 0.3, extension 20 pixels, soft edge 6 Å) and with tighter local constrains (offset_range 3, offset_step 1, sigma_ang 2) produced a 3.9 Å map. To obtain a final locally filtered and locally sharpened map, the same procedure as the one used for the microtubule datasets was used, leading to the final map (**Fig 1f, bottom structure**).

### Cryo-EM resolution estimation

The final resolutions for each cryo-EM reconstruction were estimated from FSC curves generated with Relion 3.1 post-process (FSC_0.143_ criteria, **Table 2**, **Supplementary Fig. 2**). To estimate the overall resolution, these curves were computed from the two independently refined half maps (gold standard) using soft masks that isolate a single asymmetric unit containing a kinesin and a tubulin dimer. The soft masks were created with Relion 3.1 relion_mask_create (for micro-tubule datasets: low pass filtration: 15 Å, threshold: 0.1, extension: 2 pixels, soft edge: 5-6 pixels; for the CT-*Ca*Kip3-MDC-ANP data set: low pass filtration: 20 Å, threshold: 0.1, extension: 4 pixels, soft edge: 10 pixels) applied on the correctly positioned EMAN pdb2mrc density map generated with the coordinates of the respective refined atomic models. FSC curves for the tubulin or kinesin parts of the maps were generated similarly using the corresponding subset of the PDB model to mask only a kinesin or a tubulin dimer (**Supplementary Fig. 2, Table 2**).

The final cryo-EM maps together with the corresponding half maps, masks used for resolution estimation, masks used in the partial signal subtraction for the microtubule datasets, and the FSC curves are deposited in the Electron Microscopy Data Bank (**Table 2**).

### Model building from cryo-EM densities

Atomic models of the cryo-EM density maps were built as follow. First, atomic models for the kinesin chains were generated from their amino-acid sequence by homology modeling using Modeller^61^. The initial tubulin structures were taken from^13^. Second, the protein chains were manually placed into the cryo-EM densities and fitted as rigid bodies using UCSF-Chimera^62^. Third, the models were flexibly fitted into the density maps using Rosetta for cryo-EM relax protocols^63,64^ and the models with the best scores (best match to the cryo-EM density and best molprobity scores) were selected. Fourth, the Rosetta-refined models were further refined against the cryo-EM density maps using PHENIX real space refinement tools^49^ and/or ISOLDE^65^. Fifth, the models were edited using Coot^48^. Several iterations of Phenix real space refinement and Coot editing were performed to reach the final atomic models. Atomic models and cryo-EM map figures were prepared with UCSF-Chimera^62^ and ChimeraX^29^.

## Data availability

Atomic coordinates of the *Ca*Kip3-MDN-ADP X-ray crystal structure have been deposited in the Protein Data Bank (PDB) under the accession code 7LFF (**Table 1**). Atomic coordinates and corresponding cryo-EM density maps, including the half maps, masks and FSC curves used to estimate spatial resolution have been deposited in the Protein Data Bank (PDB) and Electron Microscopy Data Bank (EMDB) under the accession codes 7TQX, 7TQY, 7TQZ, 7TR0, 7TR1, 7TR2, 7TR3 and EMD-26074, EMD-26075, EMD-26076, EMD-26077, EMD-26078, EMD-26079, EMD-26080 (**Table 2**).

## Acknowledgements

This work was supported by Canadian Institutes of Health Research Grant PJT-169149) (J.S.A.), National Sciences and Engineering Council of Canada grant RGPIN-2019-05924 (J.S.A), and National Institutes of Health Grant R01GM113164 (H.S.). Cryo-EM data collection was performed at the Simons Electron Microscopy Center and National Resource for Automated Molecular Microscopy located at the New York Structural Biology Center, supported by grants from the Simons Foundation (SF349247), NYS-TAR, and the NIH National Institute of General Medical Sciences (GM103310) with additional support from Agouron Institute (F00316) and NIH (OD019994). We thank Peter Davies for the use of microscopy facilities. We thank Huihui Kuang, Daija Bobe, Carolina Hernandez, Kashyap Maruthi, Kasahun Neselu and Mykhailo Kopylov for assistance and support during data collection. B.H. is an NSERC Canada Graduate Scholarship and Ontario Graduate Scholarship recipient.

## Author contributions

B.H., C.D. and D.T. cloned, expressed, and purified the kinesin proteins; B.H. and J.S.A collected X-ray diffraction data for the *Ca*Kip3-MDN crystal, determined the structure, built and refined the atomic model, and analyzed the structure; M.P.M.H.B. and A.B.A. assembled kinesin-microtubule and kinesin-curved tubulin complexes and made cryo-EM grid samples; A.B.A. performed sample screening and optimization for cryo-EM imaging and performed microtubule selection; M.P.M.H.B. and A.B.A. designed the cryo-EM experiments and performed cryo-EM data collection; M.P.M.H.B. designed and performed the cryo-EM data processing of kinesin-microtubule and kinesin-ring complexes, mechanistically interpreted the cryo-EM structures and proposed several of the biochemical experiments; M.P.M.H.B. and H.S. built and refined the atomic models, and analyzed the structures; B.H. designed the biochemical analyses; B.H. and C.D. performed ATPase assays; B.H. performed all other biochemical experiments; J.S.A. and H.S. supervised the project; B.H., M.P.M.H.B., A.B.A., H.S. and J.S.A wrote the manuscript.

## Supplementary Data

**Supplementary Table 1.** Extra data collection and refinement parameters.

| **Dataset** | **Microscope used** | **Pixel Size (Å)** | **Box size of symmetric map (pixels) <sup>a</sup>** | **Magnification anisotropy % <sup>b</sup>** | **Magnification anisotropy angle <sup>b</sup>** | **Box size (pixels) <sup>c</sup>** |
| --- | --- | --- | --- | --- | --- | --- |
| MT-CaKip3-MDC-ANP | Krios 3 | 0.825 | 664 | 0.54 | 48.4 | 416 |
| MT-CaKip3-MDC-AAF | Krios 2 | 0.849 | 664 | 1.50 | 178.3 | 416 |
| MT-CaKip3-MDC-APO | Krios 3 | 0.825 | 664 | 0.71 | 46.7 | 416 |
| MT-CaKip3-MDN-ANP | Krios 1 | 0.830 | 664 | 0.46 | 83.7 | 416 |
| MT-CaKip3-MDN<sub>L2-HsKHC</sub>-ANP | Krios 2 | 1.090 | 512 | 1.16 | 177.2 | 320 |
| MT-CaKip3-MDN<sub>L2-HsKHC</sub>-APO | Krios 2 | 0.849 | 664 | 1.50 | 177.7 | 416 |
| CT-CaKip3-MDC-ANP | Krios 3 | 1.083 | 700 | n.c. | n.c. | 416 |

<sup>a</sup> Box size used during the helical refinement of the microtubule datasets (MT) or during the full C14 ring refinement (CT-CaKip3-MDC-ANP). The following local refinements use a smaller box size (see Note c).
<sup>b</sup> Given when estimated and corrected for a given dataset. n.c.: magnification anisotropy not corrected.
<sup>c</sup> Box size used for the local refinements and final reconstructions.

**Supplementary Fig. 1:**
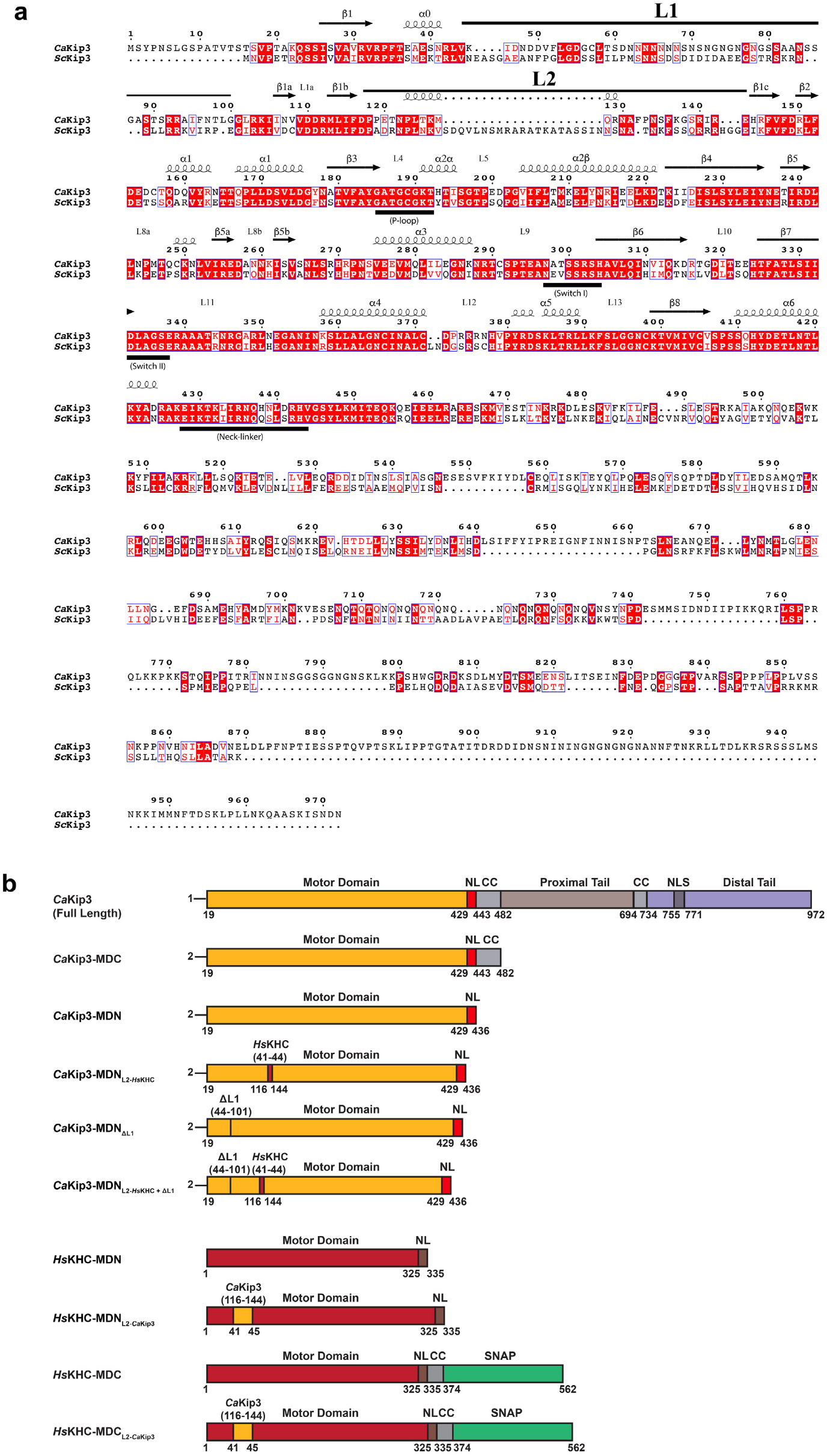
Alignment of *Ca*Kip3 and *Sc*Kip3 protein sequences and schematic of *Ca*Kip3 and *Hs*KHC constructs. (a) The numbering relates to residues of *Ca*Kip3. Identical residues are colored in white on a red background; similar residues are colored in red font. The sequence alignment was performed using Needle at the EMBL-EBI^66^. Secondary-structure elements are placed above the sequence alignment according to their positions in the crystal structures of *Ca*Kip3 (PDB entry 7LFF) using ESPript^67^. The loop-1, loop-2, the ATP-binding pocket, and the switch regions are indicated. (b) Schematic illustration of the structural motifs in *Ca*Kip3 and the truncation sites for all *Ca*Kip3 and *Hs*KHC constructs used in this study. The neck-linker (NL), predicted coiled-coil forming regions (CC), predicted nuclear localization sequence (NLS), predicted proximal and distal regions of the *Ca*Kip3 tail domain, and SNAP tag sequence are shown.

**Supplementary Fig. 2:**
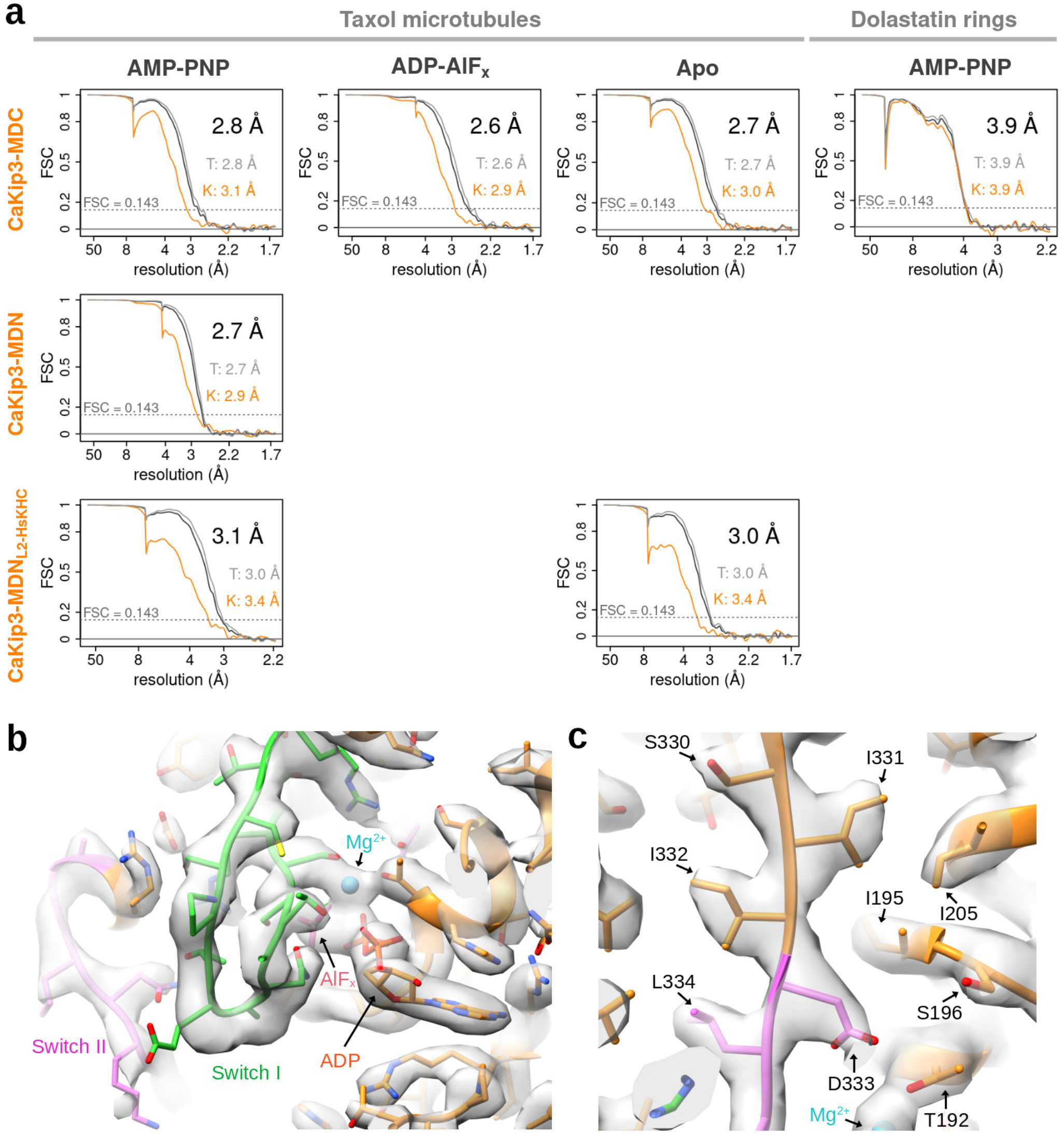
Resolution estimation of the cryo-EM maps. **(a)** FSC curves for each of the structures solved by cryo-EM. Overall FSC in black, tubulin part FSC in grey and kinesin part FSC in orange. Resolution values (FSC_0.143_) for the overall, tubulin (T) and kinesin (K) parts are indicated. Half maps and masks used to generate the FSC curves are deposited in the EMDB (accession numbers in Table 2). **(b)** Iso-density surface representation of the MT-*Ca*Kip3-MDC-AAF map with underlying model, showing the well resolved nucleotide pocket area of the kinesin in the closed state. The key elements, Switch I, Switch II, ADP, Mg^2+^ and AlF_x_, are labeled. **(c)** Inset of the MT-*Ca*Kip3-MDC-AAF map showing a strand of the central beta-sheet of the motor and neighboring region. *Ca*Kip3 residues present in this area are labeled. The figure was made with GNU R^68^ and USCF Chimera^62^.

**Supplementary Fig. 3:**
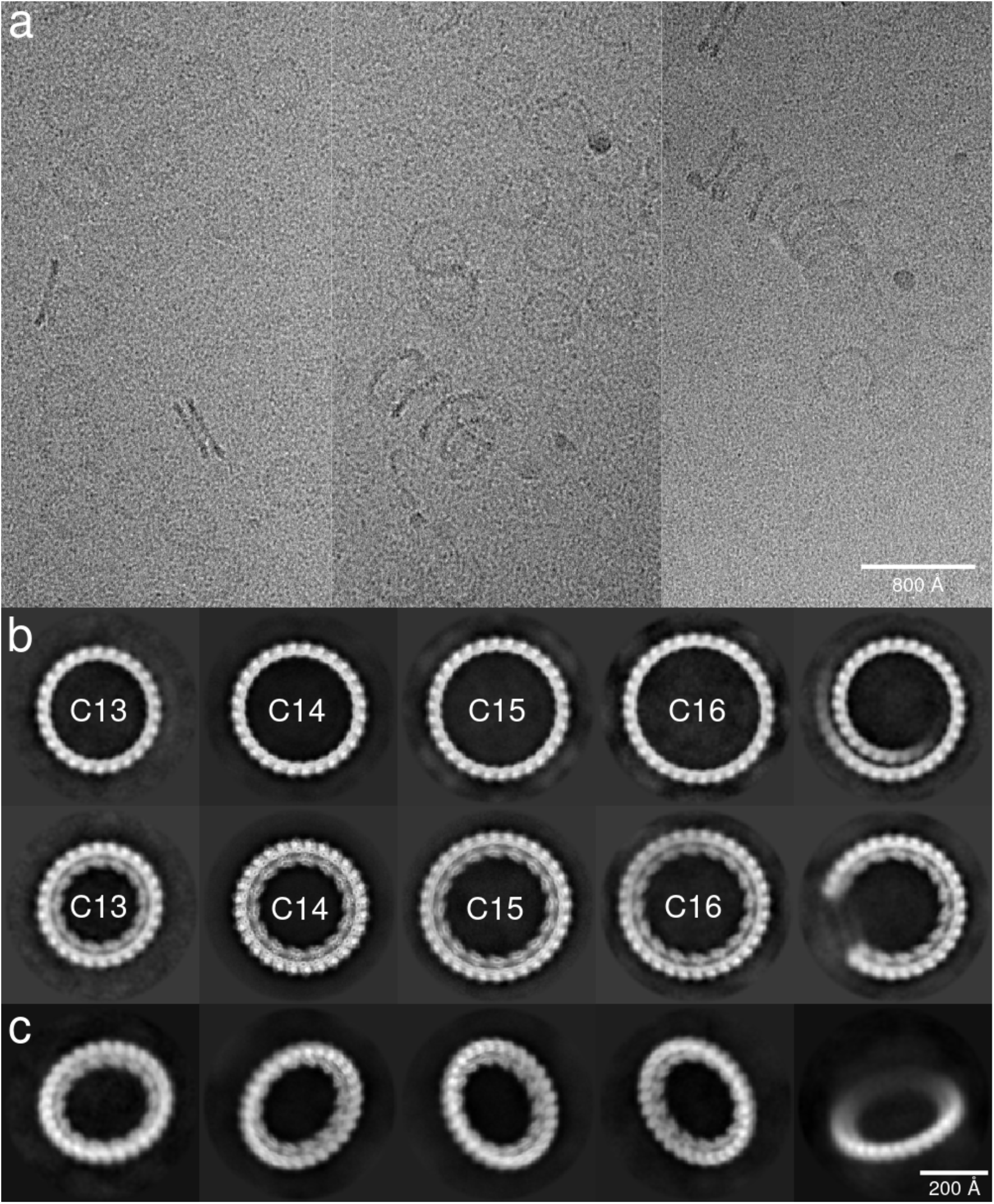
Polymer structures detected in the dolastatin-10 tubulin sample with *Ca*Kip3-MDC and 4 mM AMP-PNP. **(a)** Three micrograph subsets extracted from 3 distinct micrographs of the tilted CT-*Ca*Kip3-MDC-ANP dataset. These micrographs represent well the different curved tubulin polymer structures seen. Both rings and spiral/spring flexible structures are present. **(b)** Representative class averages of the different structures that were detected by 2D classification performed on the dolastatin-10 tubulin sample mixed with *Ca*Kip3-MDC and 4 mM AMP-PNP and collected with no stage tilt. This 2D classification was done after picking ring-like structures i.e., the numerous spring/spiral were avoided. The dataset having strong preferential orientation, the resulting 2D class averages facilitates the identification of the different structures present. The classes averages correspond to tubulin protofilament structures either undecorated (top row) or decorated with *Ca*Kip3-MDC (bottom row). This dichotomy is possibly related to a high cooperative binding behavior of *Ca*Kip3. The tubulin ring composition varies from 13 to 16 tubulin dimers; C*n* labels indicate rings with *n* tubulin dimers. The decorated C14 class is preponderant (note the more detailed class average as a consequence). The scale is the same as in panel **(c)**. **(c)** Subset of the class averages of decorated C14 rings obtained after the third cycle of autopicking/centering/2D classification done in the processing of the CT-*Ca*Kip3-MDC-ANP dataset collected with a 40-degree stage tilt. These class averages show various degrees of out of plane tilt. The figure was made with GNU R^68^.

**Supplementary Fig. 4:**
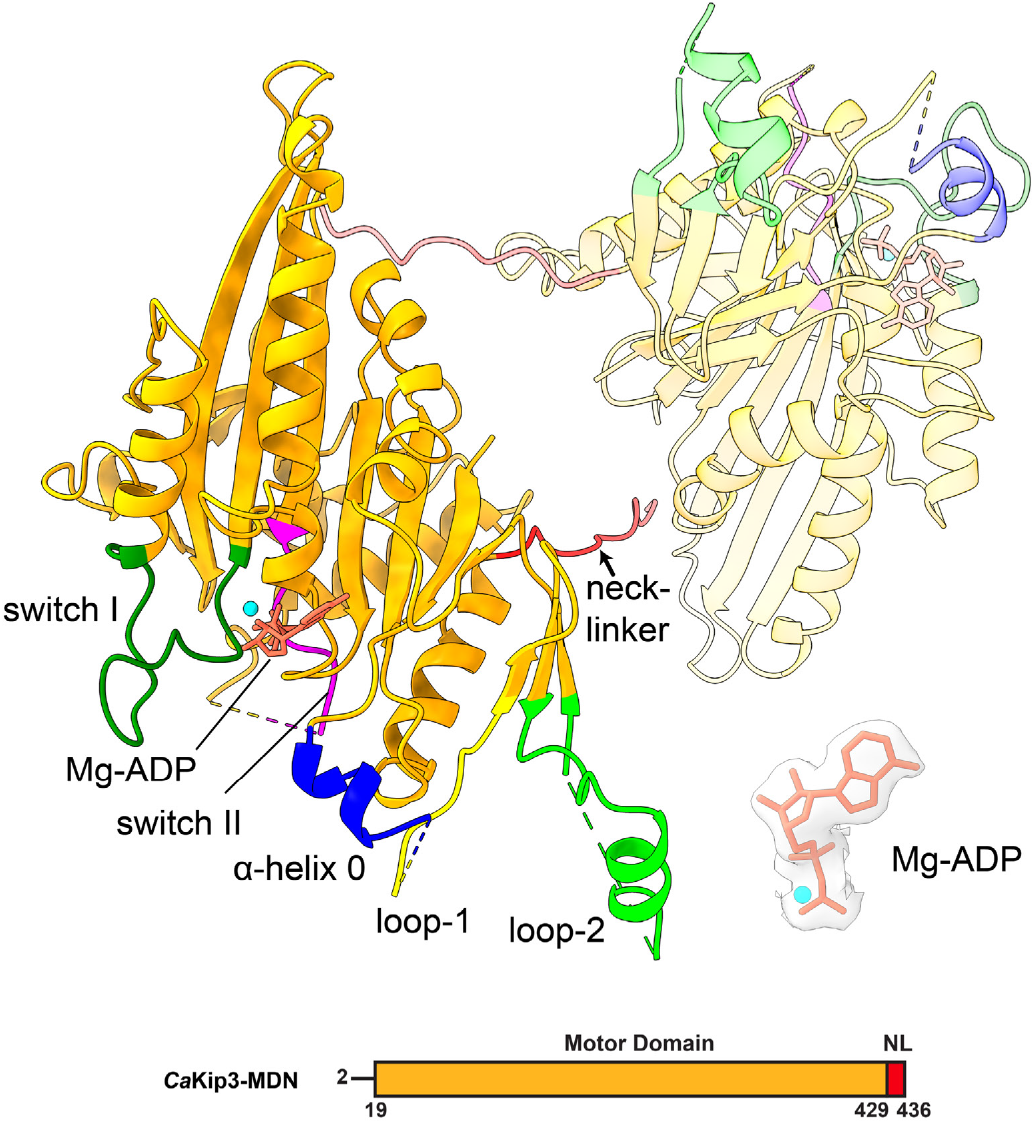
X-ray crystal structure of *Ca*Kip3-MDN. The motor core of each of the two molecules in the asymmetric unit is colored orange, one of which is shown as a transparent model. Specific regions of the motor domain that are involved in catalytic activity are colored according to **Fig. 1**. The inset panel shows the electron density for Mg-ADP from a 2m*F*_o_ − D*F*_c_ map contoured at 1.0 σ using PyMOL^69^.

**Supplementary Fig. 5:**
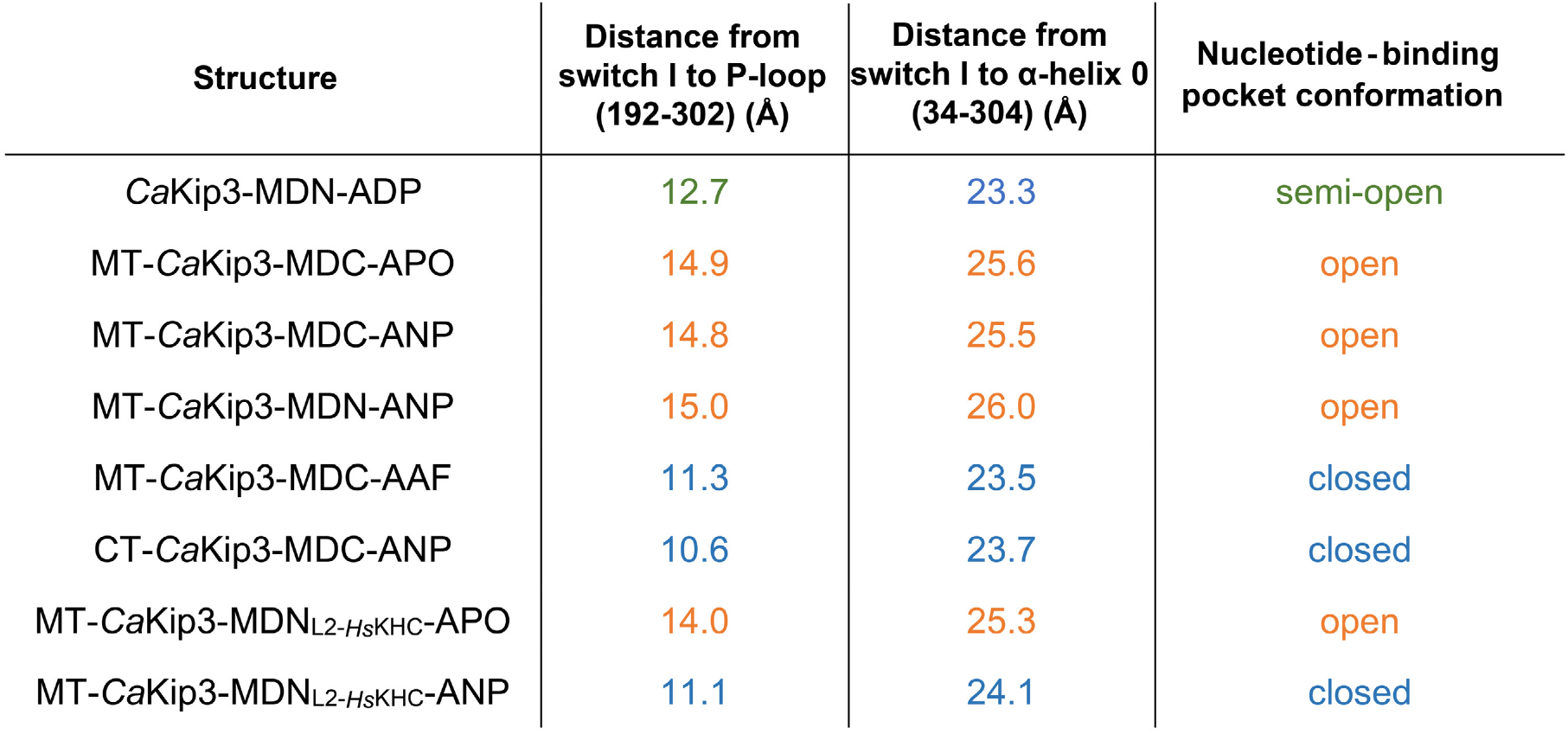
Nucleotide binding-pocket measurements for *Ca*Kip3 structures. Measurements displayed are associated with the opening and closing of the nucleotide-binding pocket^13^. Measurements were taken between the α-carbon atoms of the specified residues. Based on these distances, *Ca*Kip3 structures were categorized into three distinct groups: open (orange), closed (blue), and semi-open (green).

**Supplementary Fig. 6:**
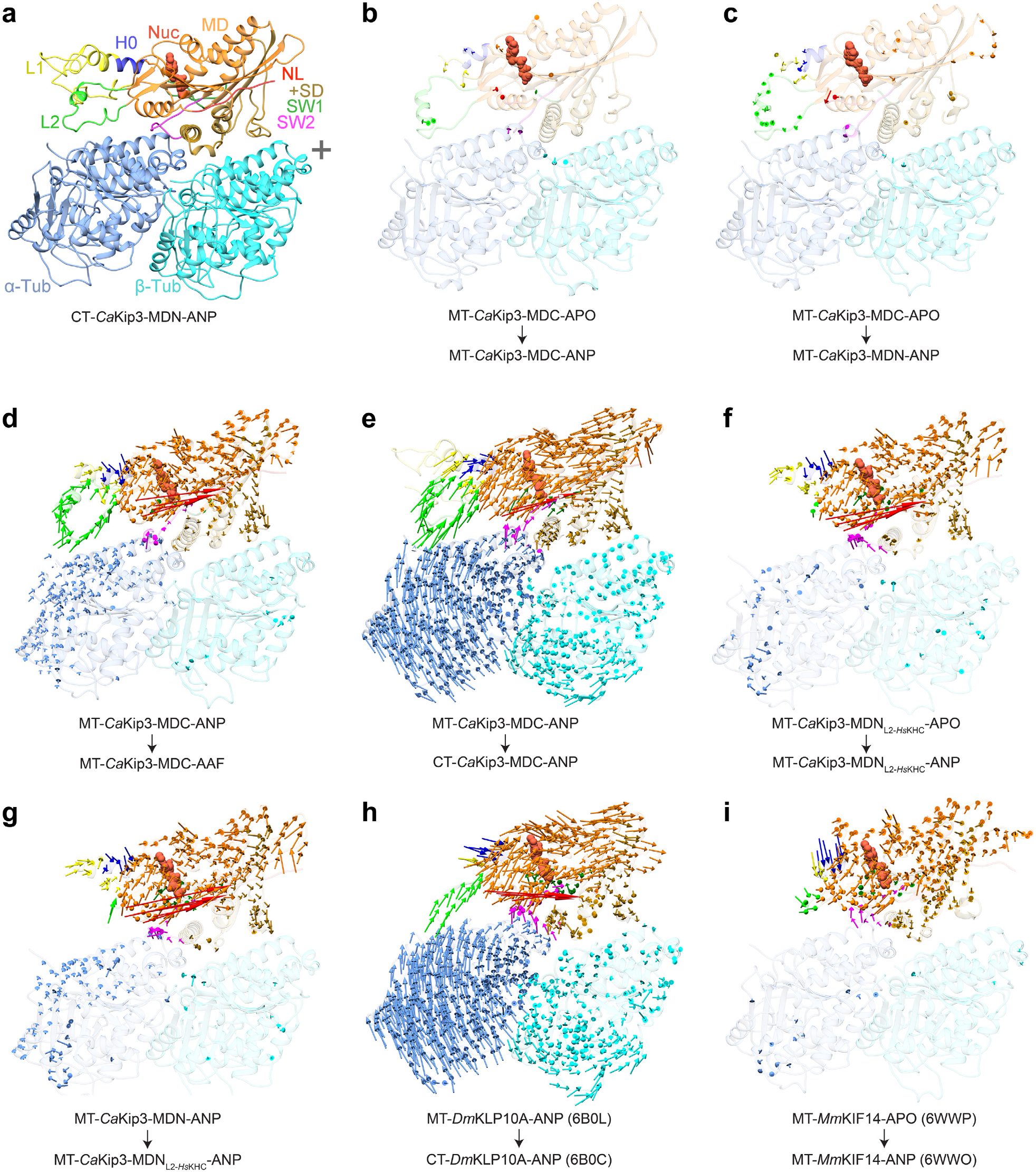
Comparison of conformational changes in kinesins and tubulin. Displacement vectors for Cα atoms in **(a-g)** *Ca*Kip3 and tubulin, **(h)** *Dm*KLP10A (kinesin-13) and tubulin, **(i)** and *Mm*KIF14 (kinesin-3) and tubulin when comparing the indicated structures. All structure comparisons were done by alignment to the β-tubulin chain. Displacement vectors for Cα atoms are colored regionally to match the segment of the protein model that is compared, using the color scheme in **Fig. 1**.

**Supplementary Fig. 7:**
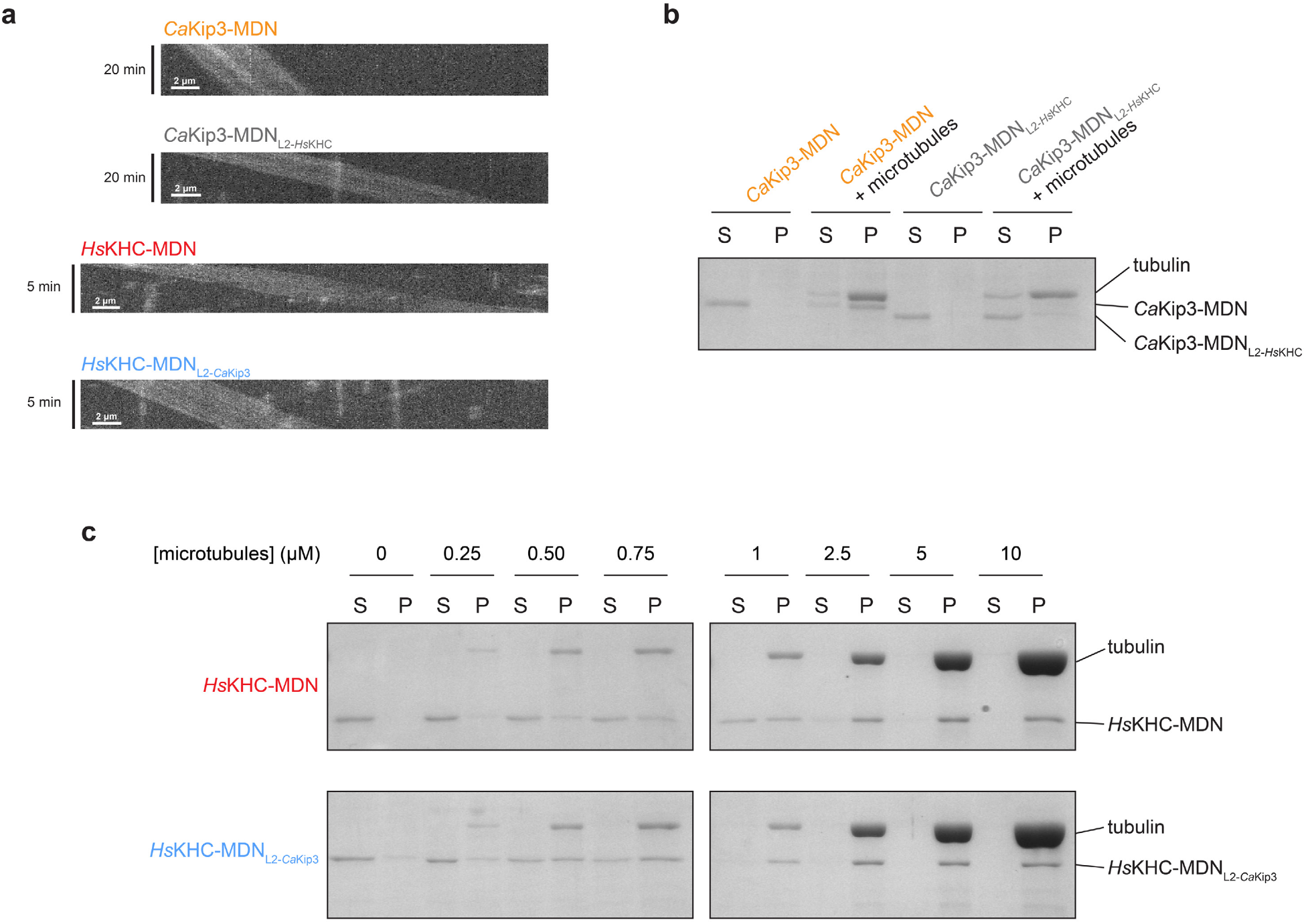
Representative data for microtubule gliding and co-sedimentation assays. **(a)** Representative kymographs from the microtubule gliding assay for *Ca*Kip3-MDN, *Ca*Kip3-MDN_L2-*Hs*KHC_, *Hs*KHC-MDN, and *Hs*KHC-MDN_L2-*Ca*Kip3_ constructs. Frames were collected every 20 seconds over a 20-minute period for kinesin-8 constructs and every 5 seconds over a 5-minute period for kinesin-1 constructs. **(b)** Representative SDS-PAGE gels showing results of the microtubule co-sedimentation assay for *Ca*Kip3-MDN and *Ca*Kip3-MDN_L2-*Hs*KHC_. Reactions contained 1 µM kinesin, 1 µM taxol-stabilized microtubules, and 2 mM AMP-PNP. Microtubules were pelleted by centrifugation to separate the free kinesin (S) and microtubule-bound kinesin (P). SDS-PAGE followed by Coomassie brilliant blue staining was used to determine the fraction of microtubule-bound kinesin. **(c)** Microtubule co-sedimentation assay results for *Hs*KHC-MDN and *Hs*KHC-MDN_L2-*Ca*Kip3_. Reactions contained 1 µM kinesin, 0-10 µM taxol-stabilized microtubules, and 2 mM AMP-PNP. Microtubules were pelleted by centrifugation to separate the free kinesin (S) and microtubule-bound kinesin (P). SDS-PAGE and Coomassie brilliant blue staining were used to determine the fraction of microtubule-bound kinesin.

**Supplementary Fig. 8:**
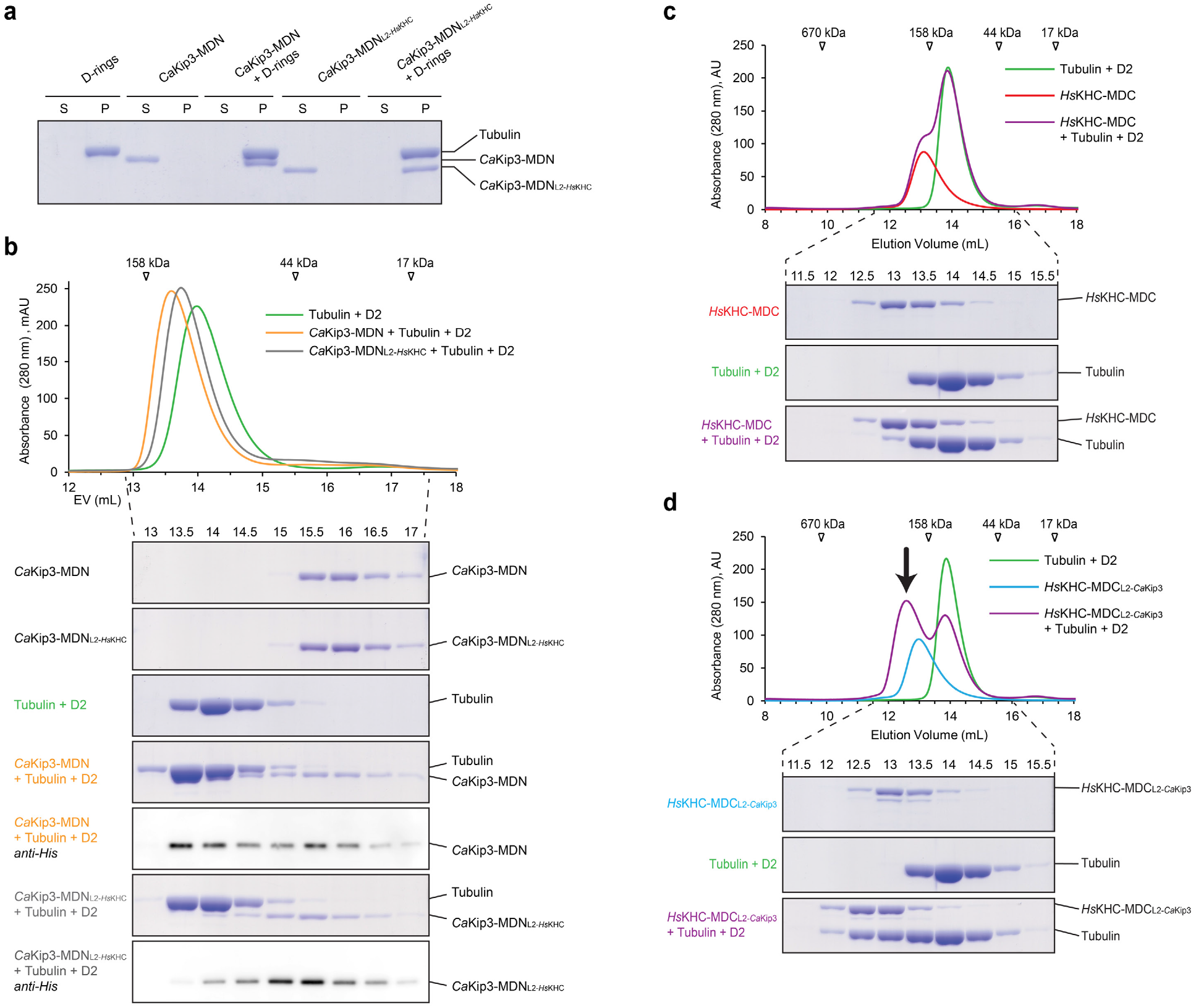
Effects of the kinesin-8 loop-2 on curved tubulin binding. **(a)** D-ring co-sedimentation binding assay performed on *Ca*Kip3-MDN and *Ca*Kip3-MDN_L2-*Hs*KHC_ at saturating conditions (4 µM kinesin, 4 µM D-rings, 20 µM dolastatin-10, 2 mM AMP-PNP). Reactions were incubated for 10 minutes, then subjected to ultracentrifugation. Supernatant (S) and pellet (P) fractions were analyzed via SDS-PAGE followed by Coomassie-blue-staining. **(b)** Size-exclusion chromatography (SEC) profiles for *Ca*Kip3-MDN and *Ca*Kip3-MDN_L2-*Hs*KHC_ in the presence of tubulin-DARPin-D2 (D2) and tubulin-D2 alone. Proteins were in a 1:1:1 molar ratio. Samples were supplemented with 0.2 mM AMP-PNP and applied to a Superdex 200 10/300 GL column in HEPES buffer. Chromatograms of *Ca*Kip3-MDN and *Ca*Kip3-MDN_L2-*Hs*KHC_ standards are omitted for clarity. SEC fractions were subjected to SDS-PAGE and stained with Coomassie blue. SEC fractions where *Ca*Kip3 could not be resolved from tubulin were additionally subjected to Western blotting analysis using anti-His antibody. **(c-d)** SEC profiles for *Hs*KHC-MDC and *Hs*KHC-MDC_L2-*Ca*Kip3_ in the presence of tubulin-D2. Black arrow on (**d**) points to the chromatogram peak observed in the *Hs*KHC-MDC_L2-*Ca*Kip3_ + tubulin + D2 sample that is not observed in the *Hs*KHC-MDC + tubulin + D2 sample. Sample preparation and column conditions same as in **(b)**.

**Supplementary Fig. 9:**
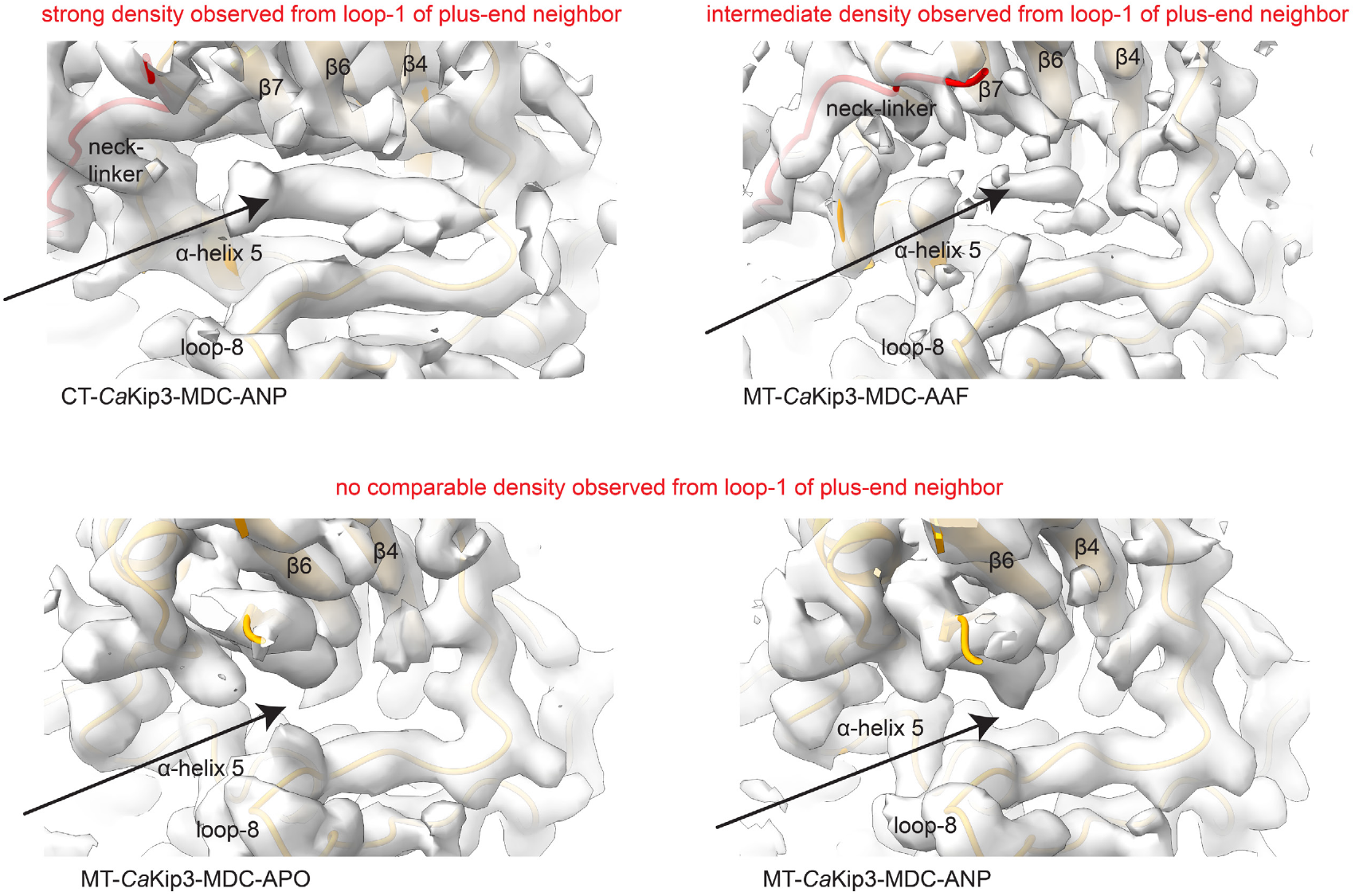
Loop-1 densities from plus-end neighboring kinesins. Densities resulting from loop-1 of the plus-end neighboring kinesin can observed strongly in the CT-*Ca*Kip3-MDC-ANP and at an intermediate intensity in the MT-*Ca*Kip3-MDC-AAF cryo-EM maps. Black arrows point to location where loop-1 densities are, or are not, observed in between the loop-8 lobe and the underside of the central β-sheet. Cryo-EM maps represented as a transparent grey surface.

**Supplementary Fig. 10:**
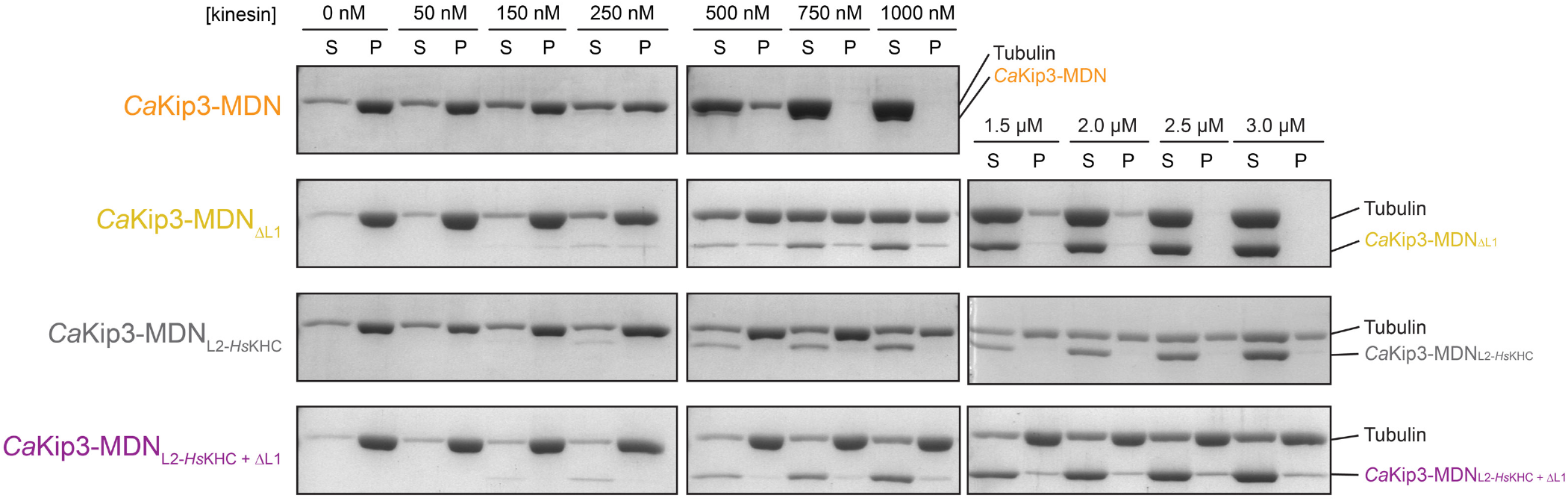
Representative data for microtubule depolymerization by sedimentation assay. Representative results are shown for *Ca*Kip3-MDN, CaKip3-MDN_ΔL1_, *Ca*Kip3-MDN_L2-*Hs*KHC_, and *Ca*Kip3-MDN_L2-*Hs*KHC+ΔL1_. Reactions were performed with 2 µM GMP-CPP-stabilized microtubules, 20 mM MgATP, and 0-3 µM kinesin in BRB80 buffer. Following a 20-minute incubation, free tubulin was separated from microtubules via ultracentrifugation. Supernatant (S) and pellet (P) fractions were subject to SDS-PAGE analysis.

**Supplementary Fig. 11:**
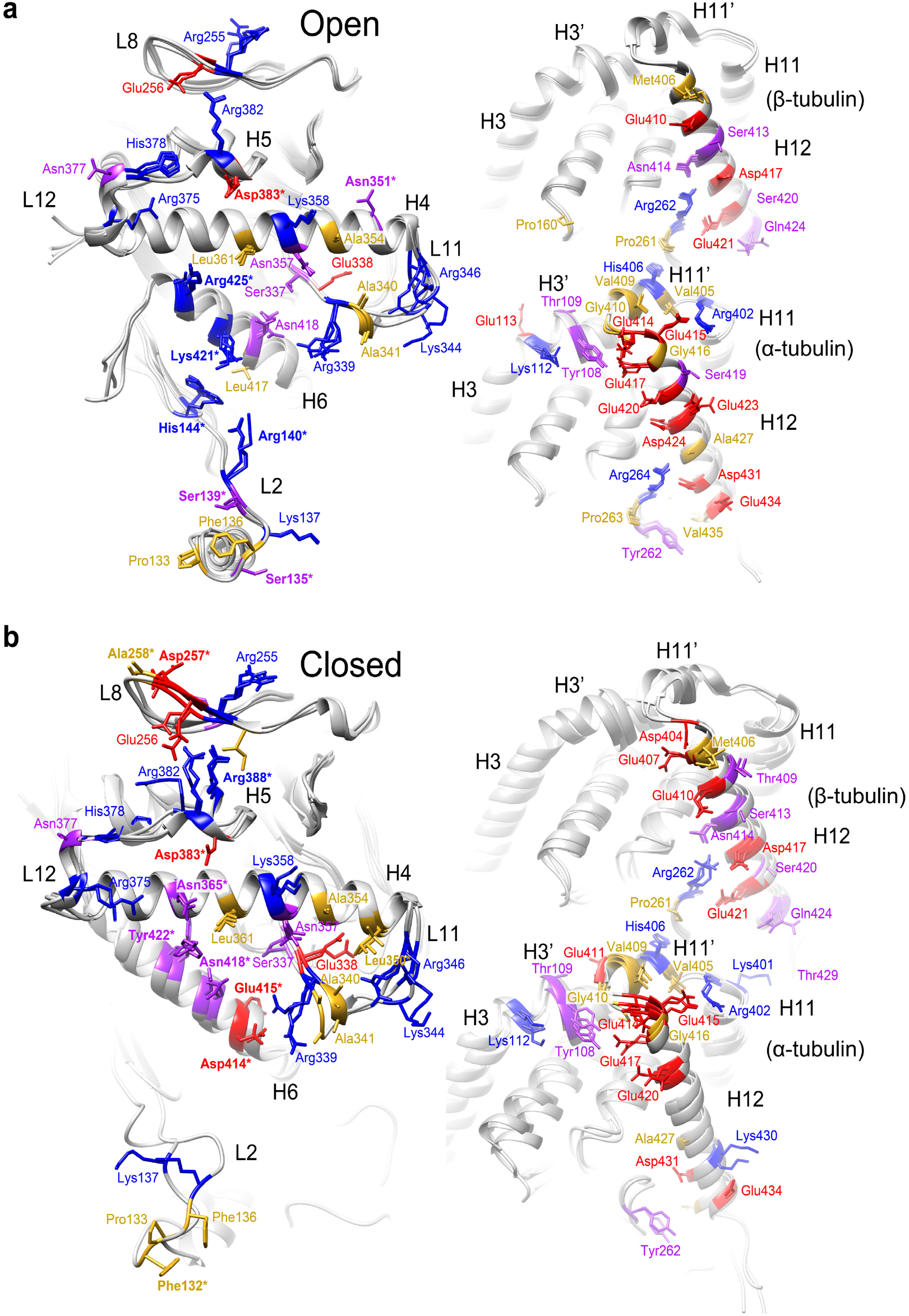
*Ca*Kip3-tubulin interacting residues. *Ca*Kip3 and tubulin residue side chains identified as making contacts by the UCSF-Chimera routine find clashes/contacts^62^. All the models exhibiting the open **(a)** or closed **(b)** conformation are superimposed. Residue side chains are colored by type (polar - purple, hydrophobic - yellow, negatively charged - red and positively charged - blue). *Ca*Kip3 residues that make contacts with tubulin in only the closed or open state are shown in bold and with an asterisk.

**Supplementary Fig. 12:**
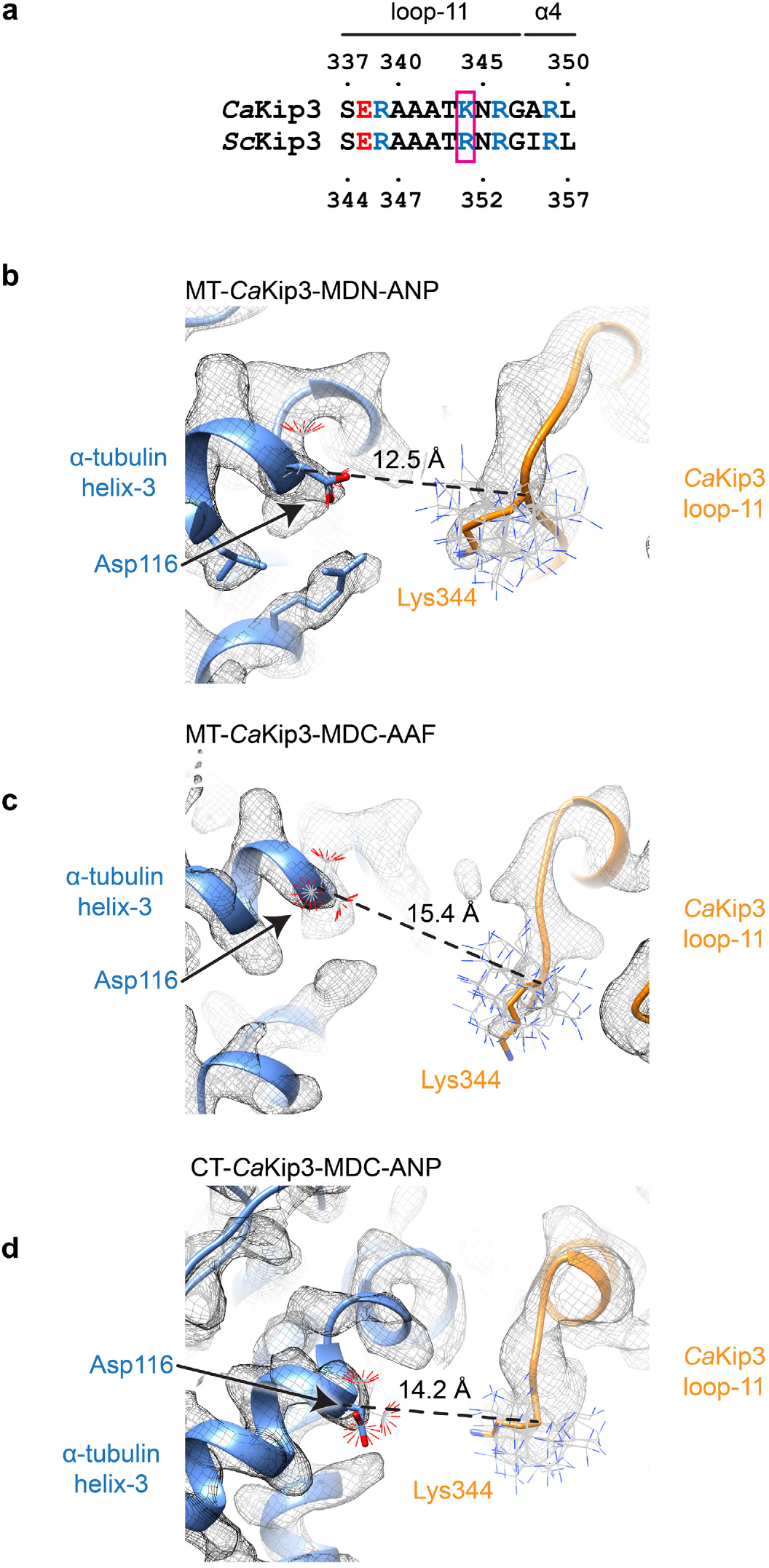
Loop-11 densities in relation to α-tubulin Asp116. **(a)** Sequence alignment of the loop-11 of *Ca*Kip3 and *Sc*Kip3. A magenta box encloses the candidate residue proposed to interact with Asp116 on α-tubulin (*Ca*Kip3 Lys344/*Sc*Kip3 Arg351)^20,21^. Close-up view of *Ca*Kip3’s loop-11 in the **(b)** MT-*Ca*Kip3-MDN-ANP, **(c)** MT-*Ca*Kip3-MDC-AAF, and **(d)** CT-*Ca*Kip3-MDC-ANP structures. The modelled conformations of *Ca*Kip3 Lys344 and α-tubulin Asp116 are shown as sticks. All possible rotamer conformations are displayed as thinner lines to depict that no side chain conformations are within bonding distance of each other. Distances displayed are from α-carbon to α-carbon. Note that on curved tubulin, the distance between Lys344 and Asp116 increases. Cryo-EM densities are displayed as a mesh surface.

